# Molecular and cellular processes disrupted in the early postnatal Down syndrome prefrontal cortex

**DOI:** 10.1101/2025.06.30.662385

**Authors:** Ryan D. Risgaard, Kalpana Hanthanan Arachchilage, Sara A. Knaack, Masoumeh Hosseini, Rachel J. Chen, Pubudu Kumarage, Danielle K. Schmidt, Xiang Huang, Jie Sheng, Carlos J. Wang, Elisa Giusti, Shuang Liu, Su-Chun Zhang, Daifeng Wang, Anita Bhattacharyya, Andre M. M. Sousa

## Abstract

Down syndrome is the most common genetic cause of intellectual disability and is characterized by early-onset delays in motor, cognitive, and language development. The molecular mechanisms underlying these neurodevelopmental impairments remain poorly understood. Here, we utilized single-nucleus multiomic sequencing to simultaneously profile gene expression and chromatin accessibility in the Down syndrome prefrontal cortex during early postnatal development, a critical period for synaptogenesis, neural maturation, and developmental neuroimmune interactions. Our findings reveal widespread dysregulation of chromatin accessibility and gene expression, with deficits spanning metabolic and synaptic pathways, oligodendrocyte lineage progression, and a pronounced neuroinflammatory signature. We present a molecular atlas of Down syndrome neuropathology at a critical stage of brain development, highlighting convergent neurodevelopmental and neurodegenerative pathways and informing potential targeted therapies for Down syndrome-associated neuroinflammation.

## Introduction

Down syndrome (DS), resulting from trisomy 21 (Ts21), is the most common genetic cause of intellectual disability^1^. Individuals with DS present in early infancy with a discernible pattern of developmental milestone delays in motor, cognitive, and language domains. This neurodevelopmental phenotype encompasses deficits in executive function, working memory, motor coordination, and linguistic processing—impairments that emerge within the first postnatal months and profoundly influence academic, occupational, and adaptive functioning trajectories throughout the lifespan^2–5^. The early onset of these deficits implicates disruptions in fundamental neurodevelopmental processes during prenatal and early postnatal corticogenesis^6^.

Despite advances in modeling Ts21, murine systems have limited translational relevance to human-specific aspects of neurodevelopment and cortical circuitry^7^, while induced pluripotent stem cell (iPSC) models have faced inherent constraints in recapitulating *in vivo* cellular diversity and the organization of neuronal-glial interactions^8,9^. To address these constraints, we performed single-nucleus gene expression and accessible chromatin (snMultiome) sequencing of the DS dorsolateral prefrontal cortex (dlPFC)—a cortical area central to higher-order cognition and working memory^10^—during the early postnatal period (0-3 years), a critical window of synaptogenesis, neuronal and glial maturation, and developmental neuroimmune interactions^11^.

Collectively, our study reveals a global dysregulation of chromatin accessibility and gene expression in the Ts21 dlPFC. This dysregulation extends beyond simple gene-dosage effects and involves complex interactions between transcription, gene regulatory networks, and the accessible chromatin landscape of discrete cell types. Our analysis reveals a pervasive dysregulation of metabolic and neuronal synaptic gene expression programs, impaired oligodendrocyte lineage progression with deficits in myelin-related transcription, and a pronounced neuroinflammatory signature marked by microglial activation and cytokine dysregulation. We reveal convergent pathways of neurodevelopmental disruption and hallmarks of incipient neurodegeneration in the early postnatal Ts21 dlPFC. This snMultiome resource (available at: https://daifengwanglab.shinyapps.io/DS_PFC/) provides a detailed, cell-type-specific molecular atlas of gene expression and chromatin accessibility in the Ts21 brain, establishing a foundation for the development of targeted therapeutic interventions addressing both developmental and degenerative aspects of Ts21 neuropathology in the early postnatal period.

## RESULTS

### Transcriptomic classification reveals selective vulnerability of cell populations in the early postnatal Ts21 dlPFC

We performed single-nucleus multiomic RNA+ATAC-sequencing (snMultiome) of the postmortem dorsolateral prefrontal cortex (dlPFC) from 5 individuals with trisomy 21 (Ts21) and 5 age- and sex-matched controls during the early postnatal period (0-3 years of age) (**Fig. 1A-B, Supplementary Table 1)**. After implementing stringent quality control measures **(Methods)**, a total of 220,956 nuclei were retained for comprehensive analysis of gene expression in the Ts21 and control dlPFC (**Fig. 1C and Supplementary Fig. 1A-J**).

**Fig. 1.**
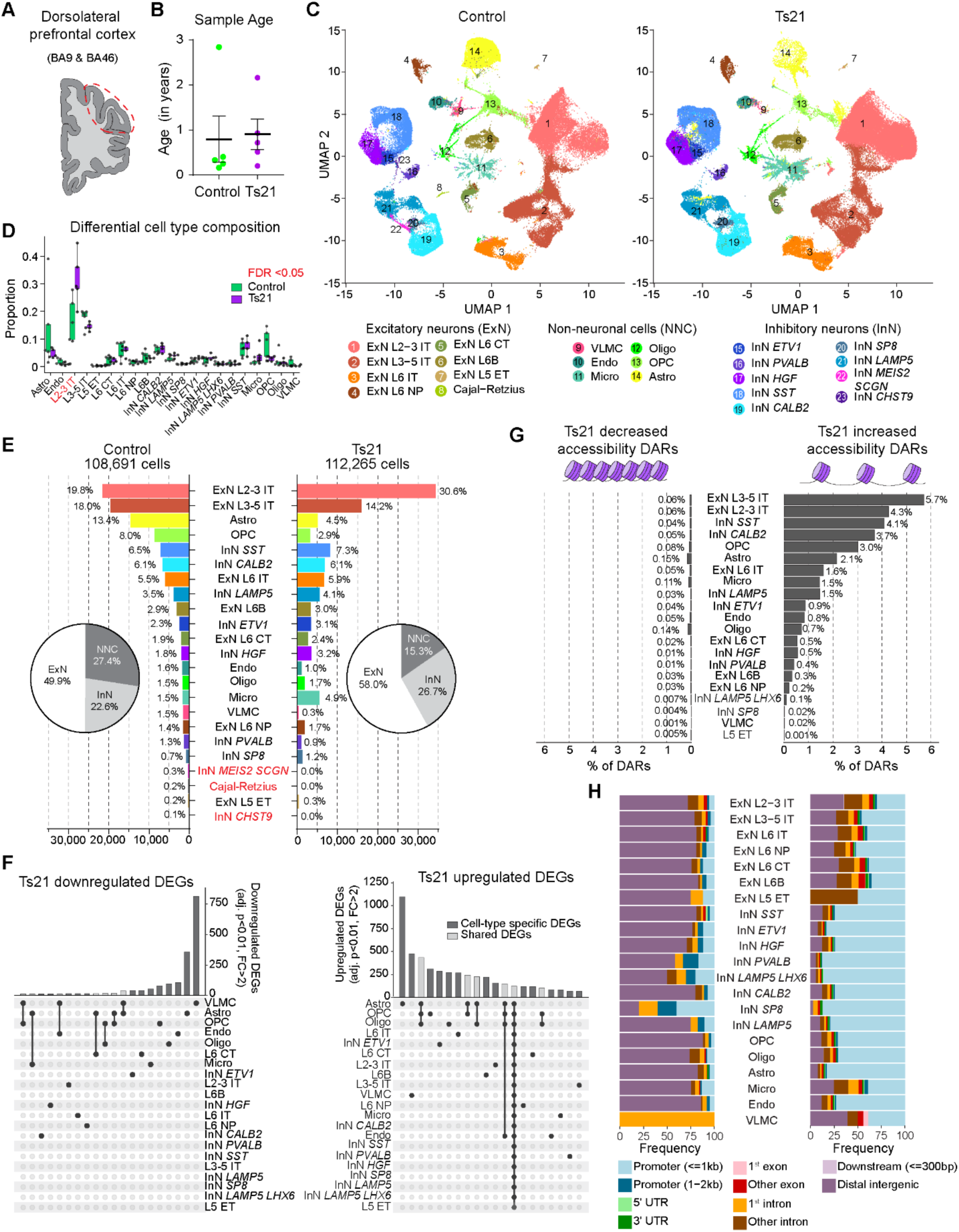
Single-nucleus multiomic gene expression and chromatin accessibility in the early postnatal Down syndrome dorsolateral prefrontal cortex (**A**) Sampling strategy of dorsolateral prefrontal cortex, encompassing Brodmann areas (BA) 9 and 46, used for single-nucleus multiomic sequencing. (**B**) Ages of samples profiled in this study. Each dot represents one biological sample. (**C**) Transcriptomic UMAP visualization of all Control (left panel) and Ts21 (right panel) nuclei analyzed in this study using a joint UMAP embedding of both Control and Ts21 nuclei, colored by cell subclass. ExN, excitatory neurons; IT, intratelencephalic; NP, near-projecting; CT, corticothalamic; ET, extratelencephalic; InN, inhibitory neurons; VLMC, vascular leptomeningeal cells; Endo, endothelial cells; Micro, microglia; OPC, oligodendrocyte precursor cells; Oligo, oligodendrocytes; Astro, astrocytes. (**D**) Differential cell type composition analysis with age and sex as covariates at the cell subclass annotation level. Statistically significant compositional changes for individual cell subclasses are highlighted in red (FDR <0.05). (**E**) Distribution and proportion of cell subclasses (bar graph) and cell classes (inset pie chart) for Control (left panel) and Ts21 (right panel). Cell subclasses present only in ontrol are highlighted in red. (**F**) Upset plot depicting differentially expressed genes at cell subclass level. Highly downregulated genes in Ts21 subclasses (adjusted p<0.01, FC>2, left panel) and highly upregulated genes in Ts21 subclasses (adjusted p<0.01, FC>2, right panel) are shown. Bars are shaded based on the cell-type specificity of the differentially expressed gene set. (**G**) Bar plot of the percentage of differentially accessible regions (DARs) per cell type with an adjusted p-value threshold of 0.05 from logistic regression, defined as the number of DARs relative to the number of peaks with observed fragments in at least 1% of barcodes for each cell type. (**H**) Genomic region annotations of DARs by cell type relative to hg38 gene annotations for Ts21 increased accessibility DARs (right panel) and Ts21 decreased accessibility DARs (left panel). See also Supplementary Figs. 1 to 3 and Supplementary Tables 1 to 4.

A fundamental question in the Down syndrome field is how an extra copy of chromosome 21 impacts the gene expression profiles both *within* and *between* cell types. To address this, we utilized an unsupervised iterative clustering approach^12,13^ to transcriptomically annotate major cell classes (excitatory neurons [ExN], inhibitory neurons [InN], non-neuronal cells [NNC]), subclasses (i.e. InN *SST*), and subtypes (i.e., InN *SST NPY CHODL*) in a hierarchical manner **(Supplementary Fig. 1K-L and Methods)**. Following batch effect correction^14^, Ts21 and control nuclei were annotated independently to minimize integration bias and identify putative Ts21-specific cell subpopulations or states. This analysis revealed a general and broad conservation of cell subclasses in the Ts21 dlPFC, with the exception of three rare (≤0.3% representation) subclasses (Cajal-Retzius cells, InN *MEIS2 SCGN,* InN *CHST9*) that were detected exclusively in control samples (**Fig. 1C** and **E, Supplementary Fig. 1K-M)**.

Next, we sought to address how Ts21 affects the identity transcriptomic cell types. To identify principal drivers of biological variability and delineate genes that maximally discriminate control and Ts21 cellular states, we performed highly variable gene (HVG) selection independently for control and Ts21, identifying the top 3000 HVGs within each condition. While the majority of HVGs were shared between control and Ts21 (71.8%), a noSupplementary Table ubset of HVGs were condition-specific **(Supplementary Fig. 2A)**. Gene Ontology (GO)^15^ analysis of control-specific HVGs revealed enrichment of homeostatic cellular processes, including gene expression, translation, and metabolism, while Ts21-specific HVGs were enriched for chronic inflammation and the response to reactive oxygen species (ROS) **(Supplementary Fig. 2B)**. Together, these findings reveal a core set of shared HVGs representing the transcriptomic variability of discrete cell types, while condition-specific HVGs suggest disrupted homeostasis, oxidative stress, and inflammation as key drivers of biological variation in the early postnatal Ts21 dlPFC.

We next sought to uncover which cell types were the most affected in Ts21. Using HVG sets calculated independently for control and Ts21, we performed unsupervised hierarchical clustering to characterize the relationship of cell subclasses between conditions **(Supplementary Fig. 2C and Methods**). This revealed approximately half (12/21) of Ts21 cell subclasses clustered based on cell identity (i.e., ExN L6B) rather than based on condition. In contrast, nine Ts21 cell subclasses demonstrated condition-specific clustering, irrespective of whether control or Ts21 HVGs were used. These subclasses predominantly comprised glial cell populations (oligodendrocyte precursor cells [OPCs], oligodendrocytes, and astrocytes), intratelencephalic-projecting (IT) excitatory neurons (L2-3 IT, L3-5 IT, and L6 IT), and caudal ganglionic eminence (CGE)-derived inhibitory neurons (InN *CALB2* and InN *SP8*) **(Supplementary Fig. 2C)**. Notably, aberrant glial and ExN populations in Ts21 clustered more closely with microglia subclasses, suggesting the activation of gene expression programs associated with inflammatory/immune responses.

To delineate transcriptomic variability *within* cell subclasses, we employed a Shannon entropy-based approach to estimate gene expression variability (**Supplementary Fig. 2D and Methods**). This demonstrated elevated transcriptomic heterogeneity in the majority (14/19) of the Ts21 subclasses analyzed. We found that all ExN subclasses exhibited greater transcriptomic heterogeneity in Ts21, while InN subclasses displayed comparative stability with subclass-specific variability. Notably, increased transcriptomic heterogeneity in Ts21 subclasses parallels observations in Alzheimer’s disease, where transcriptomic variability scores have been found to correlate with clinical severity and neurofibrillary tangle burden^16^. Collectively, these analyses identify ROS and inflammation as key drivers of biological variability in the early postnatal Ts21 dlPFC, with glial cells, IT excitatory neurons, and CGE-derived inhibitory neurons demonstrating selectively pronounced alterations in transcriptomic identity.

### Alteration of cell subclass and subtype composition in the Ts21 dlPFC

In addition to alterations in gene expression, disruptions in cellular composition and cortical arealization have been hypothesized to contribute to the pathogenesis of neurodevelopmental disorders^17,18^. To systematically evaluate compositional shifts in both broad cell subclasses and transcriptomically defined subtypes, we employed differential cell type composition analysis^19^ at both the cell-subclass and cell-subtype hierarchical levels (**Fig. 1D, Supplementary Fig. 2E-G, and Supplementary Table 2**). Cell-subclass differential abundance analysis revealed a significant increase in the proportion of upper layer excitatory neurons (ExN L2-3 IT) in Ts21 **(Fig. 1D and Supplementary Fig. 2F)**. This observation aligns with midfetal single-nucleus RNA sequencing data from our companion study^20^ while contrasting with previous findings in mouse and *in vitro* models^6^. This highlights context-dependent developmental mechanisms and the importance of analyzing human brain samples for studying human neurodevelopmental disorders.

While no significant differences were observed within InN or glial subclass proportions, subtype-level analysis revealed compositional shifts in transcriptomically-defined astrocyte, microglia, and InN subtypes (**Supplementary Fig. 2E and G**). These changes reflect likely Ts21-associated perturbations in developmental patterning, cell maturation, or subtype-specific pathological responses. Collectively, our results demonstrate a selective expansion of upper-layer IT excitatory neurons in the early postnatal Ts21 dorsolateral prefrontal cortex, concurrent with subtype-specific reorganization of glial and inhibitory neuronal populations. These laminar and cellular imbalances may contribute to the circuit-level pathophysiology observed in Ts21^21^, potentially reflecting disrupted, aberrant cortical connections and altered neuroglial interactions during the early postnatal period.

### Genome-wide transcriptional and chromatin remodeling in Ts21

To identify both shared and cell-type specific patterns of gene expression changes, we performed differential gene expression analysis^22^ at the major cell class and subclass levels (**Supplementary Table 3**). Glial cells exhibited the most pronounced upregulation of gene expression (adj. P<0.01, FC>2), whereas vascular and glial populations demonstrated the greatest number of highly downregulated differentially expressed genes (DEGs) (adj. P<0.01, FC>2) (**Supplementary Fig. 3A-B**). Although many of the Ts21 DEGs were cell-type specific, we identified a subset of highly upregulated DEGs that were present across all subclasses **(Fig. 1F)**. Notably, only 4 out of 121 of these shared upregulated genes mapped to chromosome 21, implicating secondary regulatory mechanisms—rather than direct gene dosage effects—in driving a conserved set of pathological gene expression. GO analysis of the shared Ts21-upregulated DEGs revealed significant enrichment in metabolic processes, including cellular respiration, oxidative phosphorylation, and mitochondrial function, consistent with documented mitochondrial dysregulation and bioenergetic deficits in Ts21 cells^23^ (**Supplementary Fig. 3C-D**).

The simultaneous profiling of gene expression and chromatin accessibility in single nuclei allowed us to link regulatory elements to genes, providing insights into Ts21 gene regulatory dysfunction. After implementing stringent quality control steps **(Methods)**, snATAC-seq data from 124,105 nuclei were retained for downstream analyses **(Supplementary Fig.1F-I, Supplementary Table 1)**. To assess changes in Ts21 chromatin accessibility, we calculated differentially accessible regions (DARs) at the cell subclass level **(Supplementary Table 4)**. Ts21-associated chromatin accessibility exhibited a striking asymmetry with widespread increases in accessible regions across cell types, a pattern robust to different statistical thresholds, DAR-calling methodologies, and select matched-sample analyses at the earliest study timepoint (**Fig. 1G, Supplementary Fig.3E-J, Methods)**. Ts21 DARs with increased accessibility predominantly localized to promoter regions, whereas chromatin regions with decreased accessibility were primarily confined to distal intergenic regions (**Fig. 1H**). The predominance of promoter-associated DARs suggests broad dysregulation of transcriptional activation, while distal accessibility changes reflect putative disrupted enhancer-promoter interactions. Notably, chromatin regions with increased accessibility in Ts21 demonstrated cell-type specificity in genomic region enrichment. Ts21 ExN populations had preferentially increased accessibility in intergenic and intronic regions, while InNs primarily exhibited an increase in DARs at promoters (**Fig. 1H**). Our multiomic analysis of Ts21 gene expression and chromatin accessibility revealed a global and widespread disruption of gene regulation along with discrete, cell-type-specific perturbations in the Ts21 dlPFC at a critical period of neurodevelopment. These findings underscore a global epigenetic rewiring beyond localized chromosomal effects, aligning with prior observations of genome-wide chromatin reorganization in Ts21^24–26^. Together, these results highlight the dual axis of chromatin architecture and gene expression dysregulation in Ts21, implicating both direct chromosomal and secondary regulatory effects in shaping neurodevelopmental pathology.

### Cell-cell interaction analysis reveals Ts21 dysregulated neuronal and vascular-associated signaling networks

To delineate higher-order signaling perturbations in Ts21, we constructed inferred cell-cell communication networks encompassing paracrine and autocrine signaling pathways independently for control and Ts21 (**Supplementary Table 5**)^27^. Network-level quantification of total communication probability across all cell-class pairs identified significant enrichment of Ts21 inflammatory signaling mediators, including IL-6, CCL, CXCL, SELE, and SELPLG pathways **(Supplementary Fig. 4A)**. Conversely, cell-cell adhesion (e.g., CDH1, CADM, APELIN) as well as neuronal adhesion pathways (e.g., NRXN, NCAM, NGL) were downregulated in Ts21 **(Supplementary Fig. 4A)**. At the cell-class level, we found the vast majority of Ts21 cell types displayed a greater number of paracrine and autocrine interactions than control (**Supplementary Fig. 4B**). The differential strength of these interactions, however, revealed cell-type specific patterns. Neuronal classes demonstrated a general pattern of attenuated outgoing (source) and incoming (receiver) interaction strength, whereas endothelial and vascular leptomeningeal cells (VLMCs) displayed pronounced increases in outgoing interaction strength (**Supplementary Fig. 4C**). These findings implicate cell-type specific regulatory mechanisms, with Ts21-associated suppression of synaptic adhesion gene networks in neurons contrasting with amplified vascular-associated paracrine signaling. The latter aligns with a neuroinflammatory signature, characterized by endothelial activation and cytokine pathway dysregulation, and reflective of early-stage vascular inflammatory signaling in the Ts21 brain.

### Chromosome 21 chromatin accessibility and gene expression in Ts21

Chromosome 21 (Chr21) is the genomic epicenter of Ts21 pathophysiology. To systematically evaluate Ts21-mediated genomic dysregulation, we analyzed both chromosome-specific and genome-wide perturbations in chromatin accessibility and gene expression. Consistent with well-established gene dosage effects, we found that Chr21 DEGs exhibited, on average, a ∼1.5-fold increase in expression relative to controls, with significantly higher expression levels compared to non-Chr21 DEGs across all cell subclasses (**Fig. 2A**).

**Fig. 2.**
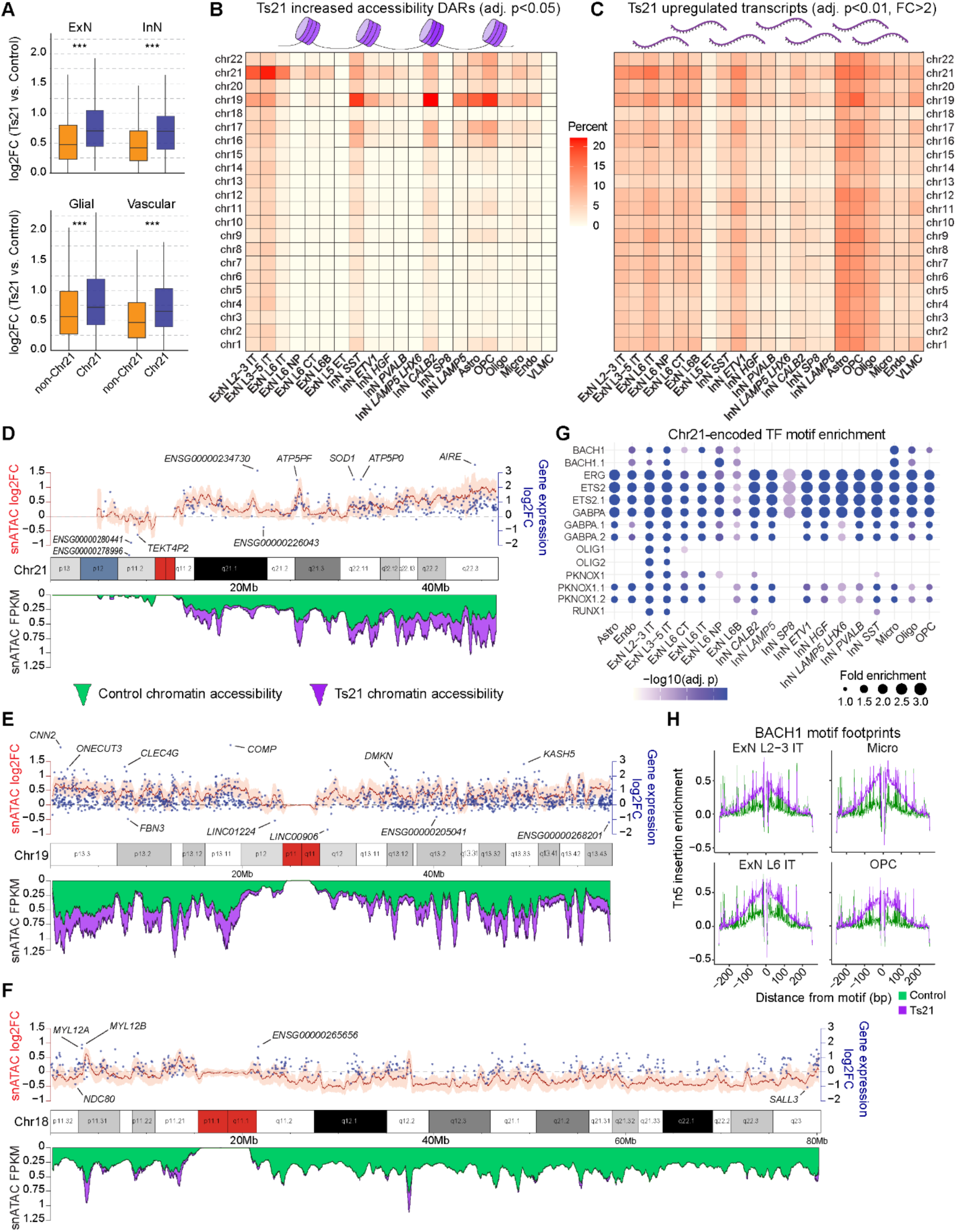
Genome-wide transcriptional and chromatin remodeling in Ts21 (**A**) Differential gene expression of chromosome 21-encoded protein-coding and lncRNA transcripts (blue) vs. non-chromosome 21 protein-coding and lncRNA transcripts (orange) across cell classes. Boxes represent the 25th and 75th percentiles and the median plotted as a central line. Whiskers represent the extension from the 25th and 75th percentile extend to the smallest and largest values within 1.5 times the interquartile range (IQR) from the quartiles and statistical significance tested using the Mann-Whitney U test. (**B**) Heatmap showing the percentage of differentially accessible regions (DARs) with increased accessibility in Ts21, grouped by cell subclass and autosome number (adjusted p<0.05). The numerator is the number of DARs, and the denominator is the number of combined peaks where a 1% fraction of barcodes showed a >0 number of fragments aligned. (**C**) Heatmap showing the percentage of highly upregulated transcripts in Ts21, grouped by cell subclass and autosome number (adjusted p<0.05, FC>2). The numerator is the number of Ts21 highly upregulated transcripts, and the denominator is the number of transcripts per chromosome. (**D to F**) Integrated visualization of gene expression and chromatin accessibility at the chromosomal scale for intratelencephalic excitatory neurons (ExN L2-3 IT, ExN L3-5 IT, ExN L6 IT) in pseudobulk. Center, ideograms for Chr21 (D), Chr 19 (E), and Chr 18 (F) depicting chromosome length in megabase pair (Mbp) and cytoband patterns. Below, snATAC FPKM coverage are depicted for control and Ts21 (green and purple, respectively) for 10 kbp bins with a rolling mean across 40 bibs. Above, the log fold change of accessibility (from FPKM units) is visualized for uniform 10 kbp bins with a rolling mean (red line) and standard deviation across 40 bins (orange shading). Mean ExN IT differential gene expression log fold change for individual genes are depicted at gene transcription start site (TSS) coordinates (blue dots). (**G)** Dot plot of Chr21-encoded transcription factor (TF) motif enrichment plotted by cell subclass (x-axis) and JASPAR motif (y-axis). Dot size represents motif fold enrichment in Ts21, and dot color represents statistical significance (adjusted p-value, hypergeometric test). (**H**) TF footprinting analysis depicting the positional distribution of Tn5 insertions within ±250 bp of TF motif of all BACH1 motif sites in peaks in select cell subclasses for control (green) and Ts21 (purple). See also Supplementary Figs. 4 and 5 and Supplementary Table 4.

To delineate genome-wide alterations in chromatin accessibility and transcription, we quantified the proportion of (1) DARs with increased accessibility and (2) highly upregulated DEGs in Ts21 across cell subclasses, stratified by chromosome (**Fig. 2B-C, Supplementary Table 4**). As expected, Chr21 displayed the most pronounced elevation in both chromatin accessibility and highly upregulated DEGs. However, this also revealed widespread non-Chr21 perturbations, with Chr19 exhibiting substantial increases in chromatin accessibility and transcriptional activity that paralleled or even exceeded Chr21-level increases for specific cell subclasses. In contrast, Chr18 demonstrated relative stability, with minimal Ts21-associated upregulation in gene expression or chromatin accessibility across cell subclasses (**Fig, 2B-C, Supplementary Fig. 4D-E**). These findings are of particular importance given the altered nuclear architecture and three-dimensional genomic organization in trisomic cells, suggesting inter-chromosomal regulatory interactions or shared mechanisms of epigenetic dysregulation at the chromosomal scale.

We next sought to examine the patterns of gene expression within chromosome 21. Given the high correlation of Chr21 gene expression within IT excitatory neuron subtypes (L2-3 IT, L3-5 IT, L6 IT) (**Supplementary Fig. 4F-G**), we grouped IT neurons in pseudobulk to examine chromatin accessibility and gene expression along the entire chromosome (**Fig. 2D)**. Interestingly, we found that Chr21 chromatin accessibility exhibited loci-specific heterogeneity. Regions spanning Chr21q22.13-q22.3 contained markedly increased chromatin accessibility and gene expression, while genic-sparse regions (Chr21q21.2) demonstrated minimal alterations in chromatin accessibility (**Fig. 2D and Supplementary Fig. 5**). This finding demonstrates the non-uniformity of chromatin accessibility and gene expression within Chr21, underscoring both localized gene dosage effects and broader trans-acting regulatory disruptions in Ts21.

Chr21-encoded transcription factors (TFs) act as pivotal regulatory nodes in Ts21 pathology, bridging localized gene-dosage effects with genome-wide transcriptional and epigenetic disruptions. To identify TFs that may be driving chromatin accessibility changes, we performed motif enrichment analysis for Chr21 TFs (**Supplementary Table 4**). This analysis identified distinct, genome-wide enrichment patterns, with members of the ETS transcription factor family (ERG, ETS2, GABPA) exhibiting pronounced overrepresentation across all cell types (**Fig. 2G**). Notably, motifs for BACH1—a TF implicated in the impairment of antioxidant defense mechanisms in Ts21^28^—were significantly enriched in accessible chromatin of excitatory neurons, microglia, and oligodendrocyte lineage cells (**Fig. 2G**). Subsequent TF footprinting analysis corroborated elevated BACH1 activity in these cell subclasses (**Fig. 2H**). Collectively, these findings underscore how the triplication of specific Chr21-encoded TFs orchestrates both cell type-specific and genome-wide regulatory disruptions, highlighting the complex relationship between chromosomal aneuploidy and global transcriptional networks.

### Metabolic and synaptic deficits in Ts21 excitatory neurons

Excitatory neurons (ExN) are the principal functional output of the dlPFC. To evaluate the effects of Ts21 on postmitotic maturing neurons, we subclustered ExNs and analyzed them independently **(Fig. 3A-B, and Supplementary Fig. 6A)**. Given our identification of pronounced transcriptomic alterations in IT excitatory neurons (**Supplementary Fig.2C-D)** and compositional alterations of upper layer *CUX2*-expressing IT neurons (**Fig. 1D and Supplementary Fig. 2E**), we focused our analyses on L2-3 IT ExNs.

**Fig. 3.**
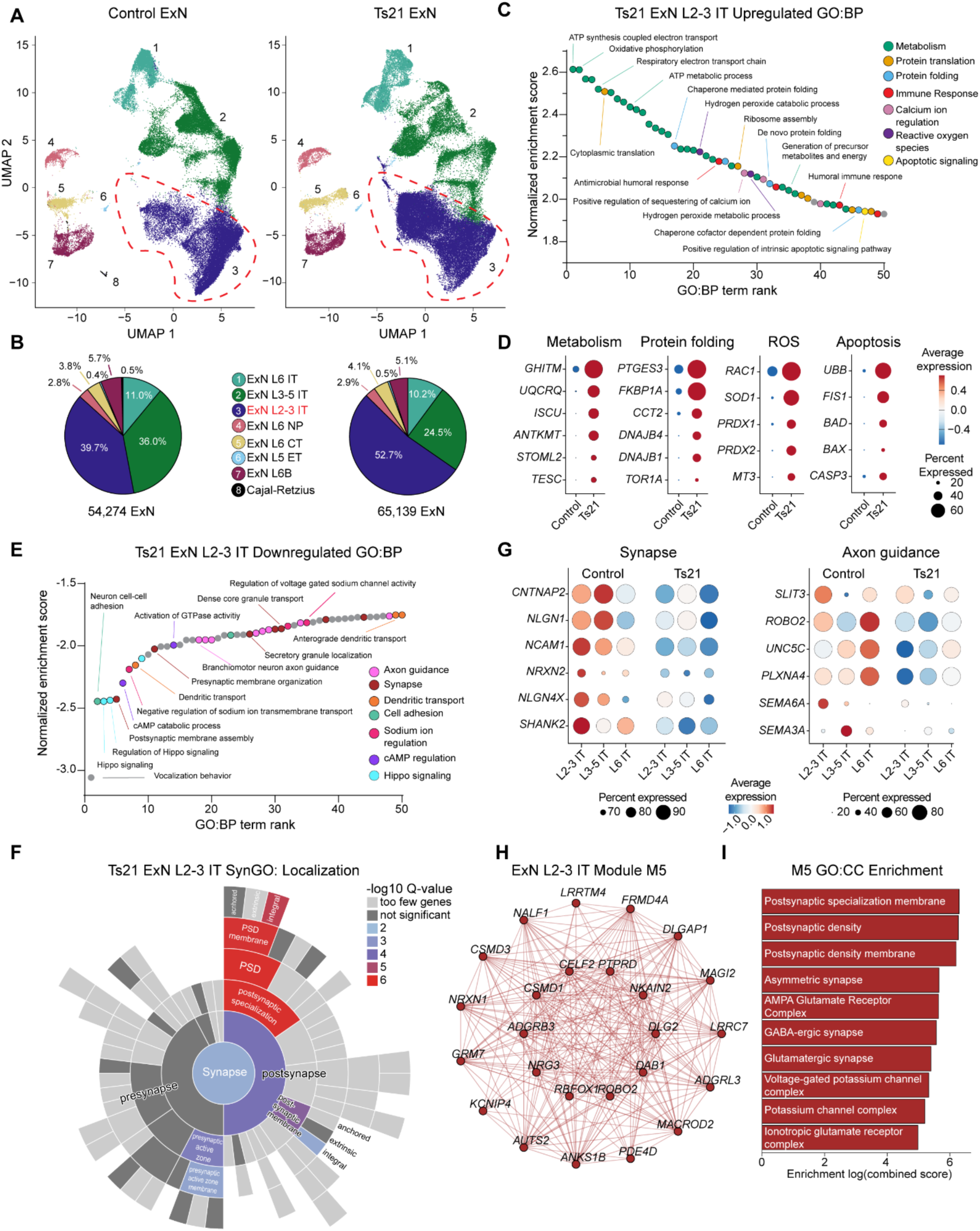
Metabolic and synaptic deficits in Ts21 intratelencephalic-projecting excitatory neurons **(A)** Transcriptomic UMAP visualization of all Control (left panel) and Ts21 (right panel) excitatory neurons (ExN) analyzed in this study using a joint UMAP embedding of both Control and Ts21, colored by cell subclass. ExN L2-3 IT subclasses are highlighted by red dashed lines. (**B**) Proportion of Control (left) and Ts21 (right) excitatory neurons colored by cell subclass. Total number of excitatory neurons analyzed in control and Ts21 is shown below the pie charts. (**C**) Waterfall plot of genes upregulated in Ts21 ExN L2-3 IT neurons (adjusted p<0.01). Gene Ontology: Biological Process (GO:BP) terms are plotted by term rank and normalized enrichment score. Select GO:BP terms are highlighted within the plot, and dots are colored by generalized biological process. (**D**) ExN L2-3 IT gene expression dot plots of select genes from generalized biological processes; size representing the percentage of nuclei expressing the gene and color representing the average expression value. (**E**) Waterfall plot of genes downregulated in Ts21 ExN L2-3 IT neurons (adjusted p<0.01). Select GO:BP terms are highlighted within the plot, and dots are colored by generalized biological process. (**F**) SynGO^30^ plot of genes downregulated in Ts21 L2-3 ExN. Each bar represents a synaptic cellular component ontology term with concentric rings representing progressively more specific subclasses of synaptic terms. Color intensity in each segment reflects the enrichment statistical significance of gene associations (-log10 Q value) for each term compared to genome-wide expectations with hypergeometric testing with Benjamini-Hochberg correction. (**G**) ExN IT subclass gene expression dot plots of select genes from synaptic and axon guidance biological processes; dot size represents the percentage of nuclei expressing the gene and color represents the average expression value. (**H**) Co-expression network visualization of Module 5 for ExN L2-3 IT neurons with the top 25 module hub genes visualized. (**I**) Bar plot depicting gene set enrichment results for ExN L2-3 IT Module 5 co-expression network with the top Gene Ontology: Cellular Components (GO:CC) enrichment terms shown. See also Supplementary Fig. 6 and Supplementary Tables 6 and 7.

Gene ontology of Ts21-upregulated DEGs in L2-3 IT ExNs **(Methods)** revealed an overrepresentation of metabolic pathways, including oxidative phosphorylation, ATP metabolism, and energy precursor generation, along with protein translation/folding and calcium ion regulation (**Fig. 3C and Supplementary Table 6**). Notably, DEGs associated with oxidative stress (ROS), immune response, and apoptotic pathways were also significantly enriched, including increased expression of pro-apoptotic regulators *BAD* and *BAX* and the executioner caspase *CASP3* (**Fig. 3D**). These findings are especially salient given the observed compositional expansion of upper-layer IT neurons in the early postnatal Ts21 prefrontal cortex (**Fig. 1D**). This is consistent with the significantly increased proportions of upper-layer *CUX2*-expressing excitatory neurons and decreased proportions of *FOXP2*-expressing L3-5 IT neurons in adolescent and adult Ts21 cohorts^29^. However, the more pronounced increase in L2-3 IT neuron composition we observed during the early postnatal period, together with elevated expression of pro-apoptotic factors, suggests prenatal overproduction of upper-layer IT identity neurons followed by apoptotic elimination during later postnatal neuronal maturation.

Ts21-downregulated DEGs in L2-3 IT neurons converged at the level of the synapse and neurite, with significant enrichment in axon guidance, synaptic organization, dendritic transport, and cell adhesion processes (**Fig. 3E**). SynGO^30^ enrichment analysis **(Methods)** identified Ts21-downregulated DEGs encoding proteins predominantly localized to the postsynaptic density (PSD) membrane, with functional roles enriched for synaptic organization (**Fig. 3F and Supplementary Table 6**). These synaptic deficits were conserved across IT subtypes, although cell-type-specific expression patterns suggest heterogeneous manifestations of a shared and widespread synaptic pathology (**Fig. 3G**). To further delineate the shared functional pathways and identify putative network-level biology, we employed high-dimensional weighted gene co-expression network analysis (hdWGCNA; **Methods**)^31^ to identify modules of Ts21 DEGs with highly correlated expression (**Supplementary Table 7**). For L2-3 IT ExN, hdWGCNA identified upregulated modules enriched for metabolic and calcium channel complexes, while the most highly downregulated module corresponded to PSD membrane components, including scaffolding proteins critical for receptor clustering and synaptic stability (**Fig. 3H-I, Supplementary Fig. 6B-F)**.

To identify cell-cell interaction patterns of synaptic and neurite deficits, we performed differential signaling analysis for all ExNs **(Methods)**. This revealed enhanced extracellular matrix-related cell signaling along with pronounced upregulation of APP signaling in Ts21 ExN (**Supplementary Fig. 6G**). We found an increased expression of *APP* across ExN subclasses, and an enhanced APP signaling network for all major cell types (**Supplementary Fig. 6H-J**). This finding is of particular significance given the chromosomal localization of *APP* on chromosome 21 and its role in the pathogenesis of amyloid beta and Alzheimer’s disease^32^. Previous work has identified significant intraneuronal amyloid-beta 42 protein accumulation in Ts21 during the early postnatal period^33,34^. This observation is pertinent, as amyloid beta has been shown to activate apoptotic pathways by disrupting calcium homeostasis and inducing the release of pro-apoptotic factors^35^, both of which are corroborated by the findings of this study.

Additionally, the most highly downregulated signaling pathways in Ts21 involved synaptic cell adhesion molecules, with many components of the neurexin, neuroligin, and neuregulin families downregulated in L2-3 IT neurons (**Supplementary Fig. 6K)**. Cell-cell interaction analyses identified these deficits are mediated, at least in part, by attenuated neuron-glia synaptic adhesion interactions (**Supplementary Fig. 6L-M**). Together, these findings reveal the molecular underpinnings for the reduced dendritic complexity, decreased synaptic density, and aberrant dendritic spine morphology that have been identified in previous Ts21 postmortem histochemical analyses^36–41^. The convergence of metabolic and ROS stress, apoptotic activation, and synaptic disorganization during a key period of synaptogenesis underscores the vulnerability of early postnatal Ts21 ExNs. Our findings link the developmental overproduction of L2-3 IT neurons with subsequent molecular, cellular, and synaptic dysfunction.

### Conserved ExN:InN neuron ratio and global synaptic dysfunction in the Ts21 dlPFC

We next sought to systematically profile the transcriptome of inhibitory neurons (InN), a cell population critical for neural circuit formation, maturation, and refinement that has been previously implicated in Ts21 pathogenesis^42^. We subclustered all inhibitory neurons and annotated them based on putative developmental origin: *SOX6* and *LHX6*-expressing medial ganglionic eminence (MGE) neurons, *PROX1* and *NR2F2*-expressing caudal ganglionic eminence (CGE) neurons, *MEIS2* and *FOXP2*-expressing dorsal lateral ganglionic eminence (dLGE) neurons, along with classical InN markers such as *SST*, *PVALB*, and *CALB2* (**Fig. 4A-B, Supplementary Fig.7A**). While mouse models of Ts21 have identified alterations in the balance of excitatory and inhibitory neurons^43–45^, we found striking conservation in the ratio of excitatory to inhibitory neurons (2.2:1) and the ratio of MGE to CGE InNs (1.4:1) in both control and Ts21 samples (**Fig. 4B**).

**Fig. 4.**
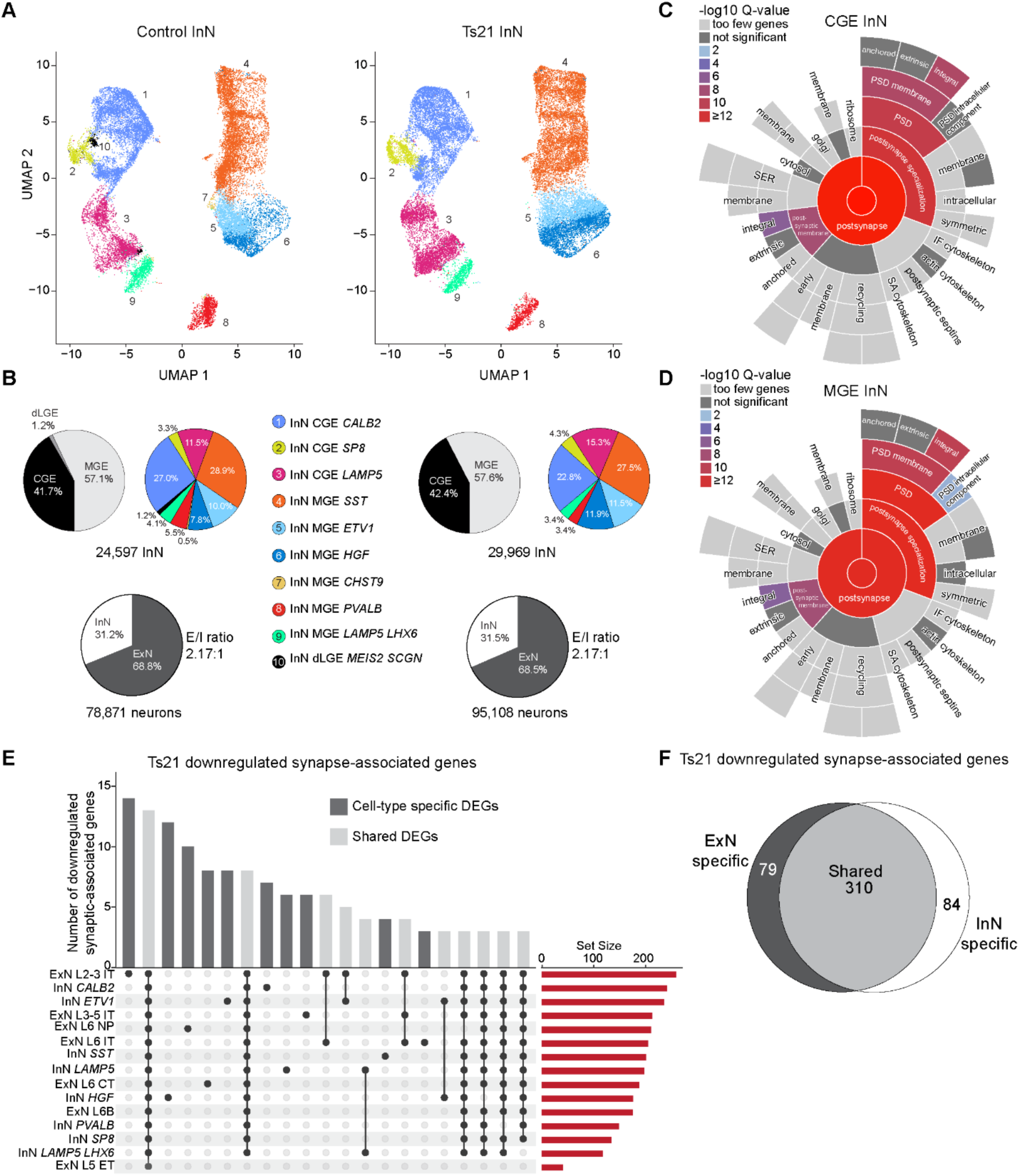
Conserved ExN:InN neuron ratio and global synaptic dysfunction in the Ts21 dlPFC (**A**) Transcriptomic UMAP visualization of all Control (left panel) and Ts21 (right panel) inhibitory neurons (InN) analyzed in this study using a joint UMAP embedding of both Control and Ts21, colored by cell subclass. (**B**) Upper pie charts: proportion of Control (left) and Ts21 (right) inhibitory neurons based on putative developmental origin or cell subclass. Lower pie charts: ratio of Control (left) and Ts21 (right) excitatory and inhibitory neurons. (**C and D**) SynGO^30^ plot of genes downregulated in CGE InN (C) or MGE InN (D). Each bar represents a synaptic cellular component ontology term with concentric rings representing progressively more specific subclasses of synaptic terms. Color intensity in each segment reflects the enrichment statistical significance of gene associations (-log10 Q value) for each term compared to genome-wide expectations with hypergeometric testing with Benjamini-Hochberg correction. (**E**) Upset of all downregulated SynGO synapse-associated genes plotted by neuronal subclass (adjusted p<0.01). (**F**) Pie chart of downregulated SynGO synapse-associated genes for ExN and InN subclasses. See also Supplementary Fig.7 and Supplementary Table 6.

As inhibitory neurons transcriptomically cluster based on their developmental origin^12^, we grouped MGE and CGE InNs and performed hdWGCNA to identify network-level perturbations in gene expression (**Supplementary Table 7**). Ts21-upregulated gene modules in MGE (MGE Module 1) and CGE (CGE Module 2) exhibited significant enrichment for metabolic and protein translation processes, whereas Ts21-downregulated modules in MGE (MGE Module 4) and CGE (CGE Module 8) were associated with PSD components, including AMPA receptor complexes and calcium channel signaling (**Supplementary Fig. 7B-K**). SynGO^30^ analysis of downregulated DEGs in both MGE and CGE InNs confirmed significant enrichment for PSD membrane-associated components (**Fig. 4C-D**). Due to this converging phenotype of downregulated genes encoding PSD proteins in inhibitory and excitatory neurons, we compared the shared and cell-type specific patterns of Ts21-downregulated synapse-associated genes.

Interestingly, we found that many synapse-associated DEGs were shared among neuronal subclasses (**Fig. 4E**). Comparative analysis of synaptic-associated DEGs revealed a high degree of overlap with 65.5% of downregulated synaptic genes shared between excitatory and inhibitory neurons (**Fig. 4F**).

These findings reveal concordant patterns of gene dysregulation among both inhibitory and excitatory neuron subclasses, uncovering a pan-neuronal molecular phenotype of metabolic and synaptic gene dysregulation. Validation via digital droplet PCR (ddPCR) with additional independent biological samples further corroborated the upregulation of ribosomal-associated genes and downregulation of synaptic genes (**Supplementary Fig. 7L-P**). Collectively, these results demonstrate that while the generation, developmental patterning, and migration of InNs to the postnatal dlPFC are generally conserved in Ts21, widespread molecular dysregulation occurs during postnatal maturation with synaptic and metabolic deficits during Ts21 interneuron circuit integration.

### Cell-type-specific regulatory deficits in Ts21 oligodendrocyte maturation and myelination

Developmental hypomyelination and oligodendrocyte deficits have emerged as defining morphological hallmarks of Ts21 pathogenesis^46^. To resolve lineage-specific perturbations, we annotated oligodendrocyte subpopulations, including *MKI67*+ proliferative oligodendrocyte precursor cells (prOPCs), *MKI67-* oligodendrocyte precursor cells (OPCs), newly formed oligodendrocytes (nfOligos), and mature myelinating oligodendrocytes (Oligos) (**Fig. 5A, Supplementary Fig. 8A, Methods)**. To investigate alterations in proliferation, differentiation, and/or maturation, we utilized pseudotime analysis^47^ coupled with PHATE^48^ dimensionality reduction to order and visualize single nuclei along a reconstructed developmental trajectory (**Fig. 5B-D and Methods**). This uncovered significantly increased pseudotime values in Ts21 OPCs and Oligos as compared to controls, with OPCs displaying the most pronounced differences, indicative of impairments in proliferation and differentiation (**Fig. 5E**).

**Fig. 5.**
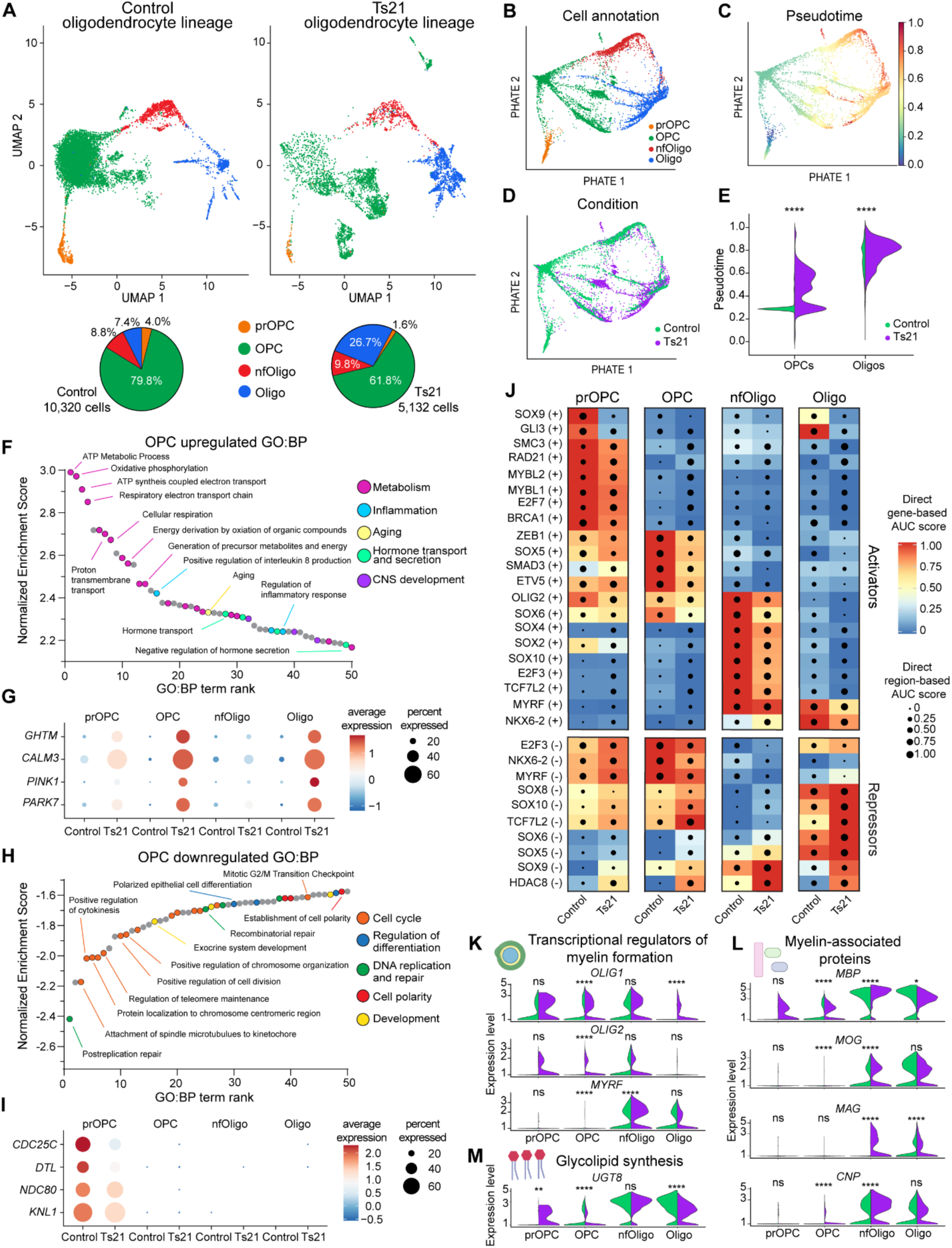
Cell-type specific regulatory deficits in Ts21 oligodendrocyte maturation and myelination (**A**) Transcriptomic UMAP visualization of all Control (left panel) and Ts21 (right panel) oligodendrocyte lineage cells analyzed in this study using a joint UMAP embedding of both Control and Ts21, colored by cell subclass. Pie charts demonstrate the proportion of each oligodendrocyte lineage cell subclass for control (left) and Ts21 (right) along with the total number of oligodendrocyte lineage cells per condition. prOPC, proliferative oligodendrocyte precursor cell; OPC, oligodendrocyte precursor cell; nfOligo, newly formed oligodendrocyte; Oligo, oligodendrocyte. (**B to D**) PHATE^48^ visualization of all oligodendrocyte lineage cells, colored by cell annotation (B), pseudotime^47^ value (C), or condition (D). (**E**) Violin plot of pseudotime^47^ values for OPCs and Oligos colored by condition (**** p<0.0001, unpaired t-test). (**F**) Waterfall plot of genes highly upregulated in Ts21 OPCs (adjusted p<0.01, FC>2). Gene Ontology: Biological Process (GO:BP) terms are plotted by term rank and normalized enrichment score. Select GO:BP terms are highlighted within the plot and dots are colored by generalized biological process. (**G**) Oligodendrocyte lineage gene expression dot plots of select genes from generalized biological processes; dot size represents the percentage of nuclei expressing the gene and color represents the average expression value. (**H**) Waterfall plot of genes highly downregulated in Ts21 OPCs (adjusted p<0.01, FC>2). Gene Ontology: Biological Process (GO:BP) terms are plotted by term rank and normalized enrichment score. Select GO:BP terms are highlighted within the plot and dots are colored by generalized biological process. (**I**) Oligodendrocyte lineage gene expression dot plots of select genes from generalized biological processes; dot size represents the percentage of nuclei expressing the gene and color represents the average expression value. (**J**) Combined heatmap and dot plot depicting select SCENIC+^51^ enhancer-driven regulon (eRegulon) activities from the oligodendrocyte lineage cell SCENIC+ gene regulatory network. Heatmap colors indicate the gene expression-based AUC scores, and dot sizes are scaled based on epigenomic region-based AUC scores. The eRegulons are grouped based on whether they are transcriptional activators (top, + regulons) or repressors (bottom, - regulons). (**K to M**) Split violin plots of genes encoding transcriptional myelin regulators (K), myelin-associated proteins (L), or glycolipid synthesis components (M) plotted by oligodendrocyte lineage cell subclass. Control and Ts21 gene expression violin plots are green and purple, respectively. (* p<0.05, ** p<0.01, *** p<0.001, **** p<0.0001, Wilcoxon Rank Sum test). See also Supplementary Fig. 8 and Supplementary Tables 6 and 8.

DEG analysis of Ts21 OPCs revealed significant enrichment for GO terms associated with metabolism, inflammation, aging, hormone transport, and development (**Fig. 5F**). Notably, we identified redox metabolism-associated genes upregulated in Ts21 OPCs and Oligos—*PINK1* and *PARK7*—that have been implicated in cellular senescence and the pathogenesis of neurodegenerative diseases^49,50^ (**Fig. 5G**). Conversely, downregulated DEGs in Ts21 OPCs were enriched for the regulation of proliferation and differentiation, including cell cycle, cell polarity, and DNA repair processes, highlighting deficits in the regulation and fidelity of mitosis (**Fig. 5H**). Indeed, kinetochore components *KNL1* and *NDC80* and key mitosis regulators *CDC25C* and *DTL* were specifically downregulated in Ts21 prOPCs (**Fig. 5I**).

To elucidate the regulatory mechanisms underlying these deficits, we constructed a single-cell multiomic gene regulatory network^51^ integrating gene expression and chromatin accessibility data from oligodendrocyte lineage cells (**Supplementary Table 8**). This analysis revealed significant attenuation of transcription factor (TF) activator activity across the Ts21 oligodendrocyte lineage, particularly affecting cell cycle-related (GLI3, E2F7, BRCA1) and oligodendrocyte lineage-specific (OLIG2, SOX10, TCF7L2, MYRF) regulons, whereas Ts21 TF repressor activity was increased in a cell-type specific manner **(Fig. 5J**). These findings uncover the gene regulatory logic and underpinnings of the observed pseudotime and differentiation impairments. Interestingly, we identified histone deacetylase (HDAC8) repressor activity as significantly increased across all oligodendrocyte lineage cell types, highlighting both shared and cell-type specific gene regulatory perturbations along the developmental trajectory (**Fig. 5J**).

The myelin sheath, a lipid-dense structure composed of phospholipids, glycolipids, and cholesterol (70%–85%) as well as myelin structural proteins (15%–30%)^52^, exhibited pronounced molecular dysregulation in Ts21 oligodendrocytes (**Supplementary Fig. 8B**). Genes encoding key cholesterol and glycolipid biosynthetic components and critical myelin-associated proteins were significantly downregulated in Ts21 (**Fig. 5K-M**). These transcriptional deficits were markedly accentuated in Ts21 mature oligodendrocytes as compared to nfOligos, revealing pathological dysregulation at the transition from newly differentiated to mature myelinating cells (**Fig. 5K-M and Supplementary Fig. 8C-I**). The transition from newly formed to myelinating oligodendrocytes was also paralleled by a pathological decline in the activity of myelin-associated gene networks. This transcriptional suppression is reflected at the gene regulatory level, evidenced by a marked decrease in the regulon activity of master TFs, specifically MYRF and NKX6-2, which are essential for myelin gene expression programs in mature oligodendrocytes^53–55^ (**Fig. 5J**).

Impaired maturation of Ts21 Oligos is likely to be mediated, at least in part, by non-cell autonomous mechanisms as evidenced by the presence of inflammation-associated DEGs across the Ts21 oligodendrocyte lineage (**Fig. 5F and Supplementary Fig. 3B-C**). Cell-cell interaction profiling revealed further attenuation of Ts21 oligodendrocyte-neuronal adhesion signaling with marked reductions in NCAM, NRXN, ADGRL, and CADM interactions, indicative of compromised axon-glial adhesion interactions (**Supplementary Fig. 8J-R**). Collectively, these results delineate a state-specific suppression of transcriptional networks governing myelination, driven by impaired regulon activity, and downstream functional impairments of myelin biosynthesis and oligodendrocyte-axonal adhesion. Together, our findings implicate dysregulated oligodendrocyte lineage progression and maturation-dependent repression of myelin gene programs as central mechanisms contributing to Ts21-associated hypomyelination, highlighting critical nodes of vulnerability in myelin gene regulatory networks.

### Molecular signatures of Ts21 astrocyte dysfunction reveal regulators of reactivity, stress response, and cell survival

Accumulating evidence has underscored the critical role of astrocytes in the pathophysiology of both Down syndrome^56^ and Alzheimer’s disease^57^. Using the pan-astrocyte marker *AQP4*, we identified and subset astrocytes from Ts21 and control. Dimensionality reduction and unsupervised hierarchical clustering demonstrated unique clustering of Ts21 astrocytes, implying fundamental alterations in cellular states or disease-associated gene expression (**Fig. 6A**).

**Fig. 6.**
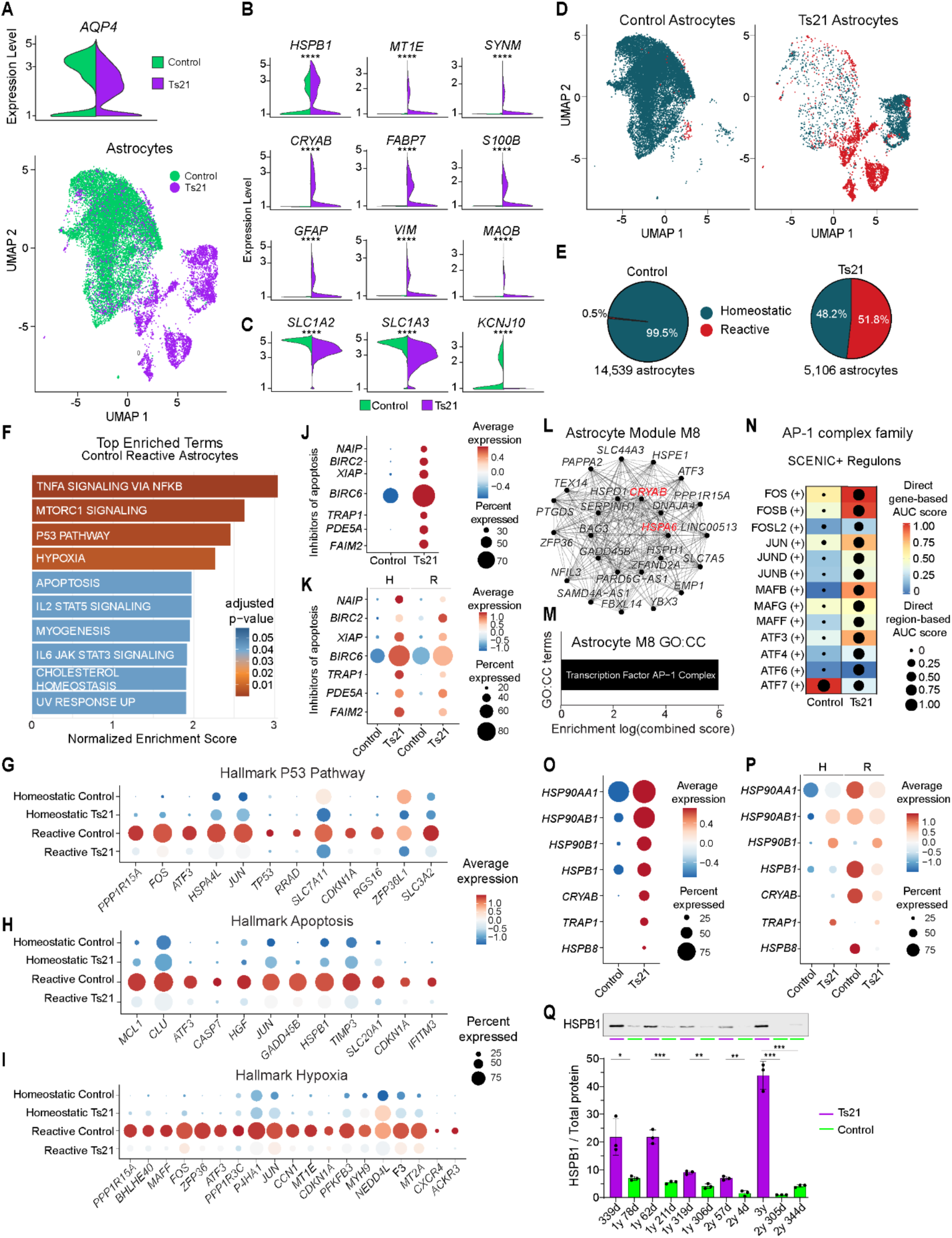
Molecular signatures of Ts21 astrocyte dysfunction reveal regulators of reactivity, stress response, and cell survival (**A**) Upper panel: astrocytes were identified and subset by expression of the pan-astrocyte marker *AQP4*. Bottom panel: Transcriptomic UMAP visualization of all Control and Ts21 astrocytes analyzed in this study using a joint UMAP embedding, colored by condition. (**B and C**) Split gene expression violin plots of control (green) and Ts21 (purple) astrocytes for consensus genes upregulated (B) or downregulated (C) in reactive astrocytes^58^. (**D**) Transcriptomic UMAP visualization of all Control (left) and Ts21 (right) astrocytes analyzed in this study using a joint UMAP embedding, colored by astrocyte state. (**E**) Proportion of control (left) and Ts21 (right) homeostatic and reactive astrocytes. (**F**) Hallmark gene set^75^ enrichment plot for genes upregulated in control reactive astrocytes as compared to Ts21 reactive astrocytes (DEGs: adj. p<0.01, Wilcoxon Rank Sum Test). Normalized enrichment score is plotted as bar height and bar color represents hallmark gene set enrichment adjusted p value (Benjamini-Hochberg procedure). (**G to I**) Astrocyte gene expression dot plots of select genes from Hallmark gene sets, (P53 pathway (G), Apoptosis (H), Hypoxia (I)) significantly enriched in control reactive astrocytes; dot size represents the percentage of nuclei expressing the gene and color represents the average expression value. **(J and K**) Astrocyte gene expression dot plots of apoptosis inhibitors stratified by condition (J) or condition and astrocyte state (K). H, homeostatic astrocytes; R, reactive astrocytes. **(L)** Co-expression network visualization of astrocyte Module 8 with the top 25 module hub genes depicted. Hub genes highlighted in red are reactive astrocyte marker genes with increased expression in Ts21 that have been validated with ddPCR in this study (Supplementary Fig.9) **(M)** Bar plot depicting gene set enrichment results for Module 8 of the astrocyte co-expression network with the top Gene Ontology: Cellular Components (GO:CC) enrichment terms visualized. **(N)** Combined heatmap and dot plot depicting AP-1 complex family SCENIC+ enhancer-driven regulon (eRegulon) activities in astrocytes. The SCENIC+^51^ gene regulatory network was obtained using all non-neuronal cells. Heatmap colors indicate the gene expression-based AUC scores and dot sizes are scaled based on epigenomic region-based AUC scores. All depicted eRegulons are transcriptional activators (+ regulons). (**O and P**) Astrocyte gene expression dot plots of cellular stress-related proteins stratified by condition (O) or condition and astrocyte state (P). H, homeostatic astrocytes; R, reactive astrocytes. (**Q**) Western blotting for HSPB1. Bar graph demonstrates densitometric analysis of HSPB1 band intensity normalized to Revert™ 700 Total Protein Stain (LICORbio). Individual dots represent blotting technical replicates (* p<0.05, **p<0.01, *** p<0.001, **** p<0.0001, two-tailed unpaired t-test). See also Supplementary Figs. 9 and 10 and Supplementary Tables 6 to 8.

Transcriptomic analysis revealed pronounced molecular perturbations in Ts21 astrocytes, including significant upregulation of reactivity-associated markers^58^ and concurrent downregulation of astrocytic homeostatic genes (**Fig. 6B-C, Supplementary Fig. 9A)**. We confirmed the expression of these upregulated reactive astrocyte markers and attenuated homeostatic genes with ddPCR validation in biologically independent, age-matched samples (**Supplementary Fig. 9C-H**). Strikingly, over half (51.8%) of Ts21 astrocytes displayed reactive molecular features in the early postnatal period compared to only 0.5% in controls (**Fig. 6D-E and Supplementary Fig. 9A**). Additionally, we found a general pattern of increasing astrocyte reactivity throughout postnatal development with ∼80% of Ts21 astrocytes displaying reactivity by two years of age, indicative of an exacerbation of astrocyte reactivity over time **(Supplementary Fig. 9B)**.

Following the identification of reactive astrocytes in both control and trisomy 21 (Ts21) samples, we sought to elucidate the conserved and divergent gene expression programs governing reactivity. DEG analysis revealed enrichment for P53 pathway, hypoxia, and apoptosis hallmark gene sets in control reactive astrocytes as compared to Ts21 reactive astrocytes (**Fig**. **6F**). Notably, we identified enrichment of *TP53* and executioner caspase *CASP7* expression specifically in control reactive astrocytes, suggesting that astrocyte reactivity in control samples is likely related to the postmortem interval, with enrichment for hypoxia-dependent gene expression and upregulation of the TP53-dependent apoptotic pathway (**Fig. 6G-I**). In contrast, Ts21 astrocytes— in both homeostatic and reactive states—displayed molecular signatures consistent with apoptosis resistance, including upregulation of apoptotic inhibitors such as *TRAP1*, *BIRC2*, and *BIRC6* (**Fig. 6J-K)**. Together, these findings uncover the recruitment of astrocytic gene expression programs for the inhibition of apoptosis and adaptive response to Ts21 pathology in the developing dlPFC.

To delineate co-expressed gene networks of astrocyte pathology, we performed hdWGCNA on Ts21 astrocyte DEGs (**Supplementary Fig. 9I-L)**. This revealed the downregulation of key genes associated with astrocyte homeostasis – *DIO2*, *SLC5A3*, and *AQP4* (Module 7), along with upregulation of a gene module consisting of established reactive astrocyte markers - *CRYAB* and *HSPA6* – and numerous heat shock protein and molecular chaperone genes (Module 8) (**Fig. 6L and Supplementary Fig. 9L**). Interestingly, this module was highly enriched for targets of the transcription factor AP-1 complex, a dimeric protein assembly critical for immune regulation and a recently identified Ts21 therapeutic target^25^ (**Fig. 6M**).

Given the drastic transcriptomic and chromatin accessibility changes in Ts21 glia, we generated a multiomic gene regulatory network encompassing all non-neuronal cells analyzed in our dataset to uncover the regulatory architecture underlying these deficits **(Supplementary Table 8 and Supplementary Fig. 10)**. This analysis revealed a marked increase in the regulon activity of canonical transcription factors governing astrocyte reactivity (STAT3, ETS2, NKFB1) and pronounced activation of AP-1 family members in Ts21 astrocytes, except for ATF7, which exhibited reduced activity (**Fig. 6N**). A Ts21-specific reduction in ATF7 regulon activity is notable, given the role of ATF7 in epigenetic repression of inflammatory pathways and inhibition of NF-Kβ signaling^59^, aligning with exacerbated inflammatory signaling in Ts21 astrocytes. Our findings reveal the Ts21 AP-1 complex as a key mediator of reactivity with TF-target gene links to dysregulated networks of astrocyte gene expression (**Supplementary Fig. 9M**). This includes downstream targets of reactivity and inflammation, most notably *JAK3*, *CRYAB*, and *APOD* (**Supplementary Fig. 9M**).

Furthermore, AP-1 targets in Ts21 astrocytes were enriched for components of cellular stress responses, particularly heat shock proteins (HSPs) and molecular chaperones. Ts21 astrocytes exhibited a broad upregulation of stress-responsive HSPs, of which we identified both Ts21-specific (*HSP90B1*) and reactivity-associated (*HSP90AA1*) enrichment (**Fig. 6O-P**). Stress response activation reflects astrocytic plasticity in Ts21, balancing pro-survival adaptations with stress-related transcriptional programs. Of particular interest was the prominent enrichment of *HSPB1*, a key stress response protein and reactive astrocyte marker that has recently been shown to be secreted by astrocytes proximal to amyloid plaques where it exerts non-cell-autonomous neuroprotection through the stabilization of proteostasis^60^. We confirmed significantly increased protein expression of HSPB1 in the early postnatal Ts21 dlPFC via immunoblotting (**Fig. 6Q**).

Collectively, these findings demonstrate that Ts21 astrocytes exhibit profound and progressive reactivity, driven by AP-1 factor complex regulation and stress response-mediated adaptation. The identification of AP-1 and stress response components as central regulators of astrocyte pathology highlights their potential as therapeutic targets for modulating astrocyte dysfunction in Down syndrome and related neurodegenerative disorders.

### Cell intrinsic gene regulation in Ts21 neuroinflammatory microglial activation

Microglia, the resident immune cells of the brain, play a critical role in activity-dependent synaptic pruning and the refinement of neural circuits during key periods of postnatal synaptogenesis^61^. To systematically analyze microglia, we annotated and subset these cells using the pan-microglial markers *APBB1IP* and *P2RY12* (**Fig. 7A**). Importantly, both control and Ts21 cells were enriched for microglia-specific markers and lacked molecular features of peripheral macrophages or other non-resident immune cells (**Supplementary Fig.11A**). In concert with our findings in astrocytes, Ts21 microglia demonstrated marked upregulation of known activated microglia markers (*ITGAX*, *SOCS3*, *FCGR2A*) alongside a significant reduction in the expression of homoeostatic genes (*SORL1*, *P2RY12*, *CX3CR1*) (**Fig. 7B-C**). Classification of microglial states based on these key markers revealed a striking shift in Ts21 microglia toward an activated phenotype, with all analyzed cells exhibiting molecular features of activation (**Fig. 7D-E and Supplementary Fig. 11B-E**). These findings were further validated using ddPCR in biologically independent and age-matched samples, revealing marked neuroinflammation in the early postnatal Ts21 dlPFC (**Supplementary Fig. 11F-J**).

**Fig. 7.**
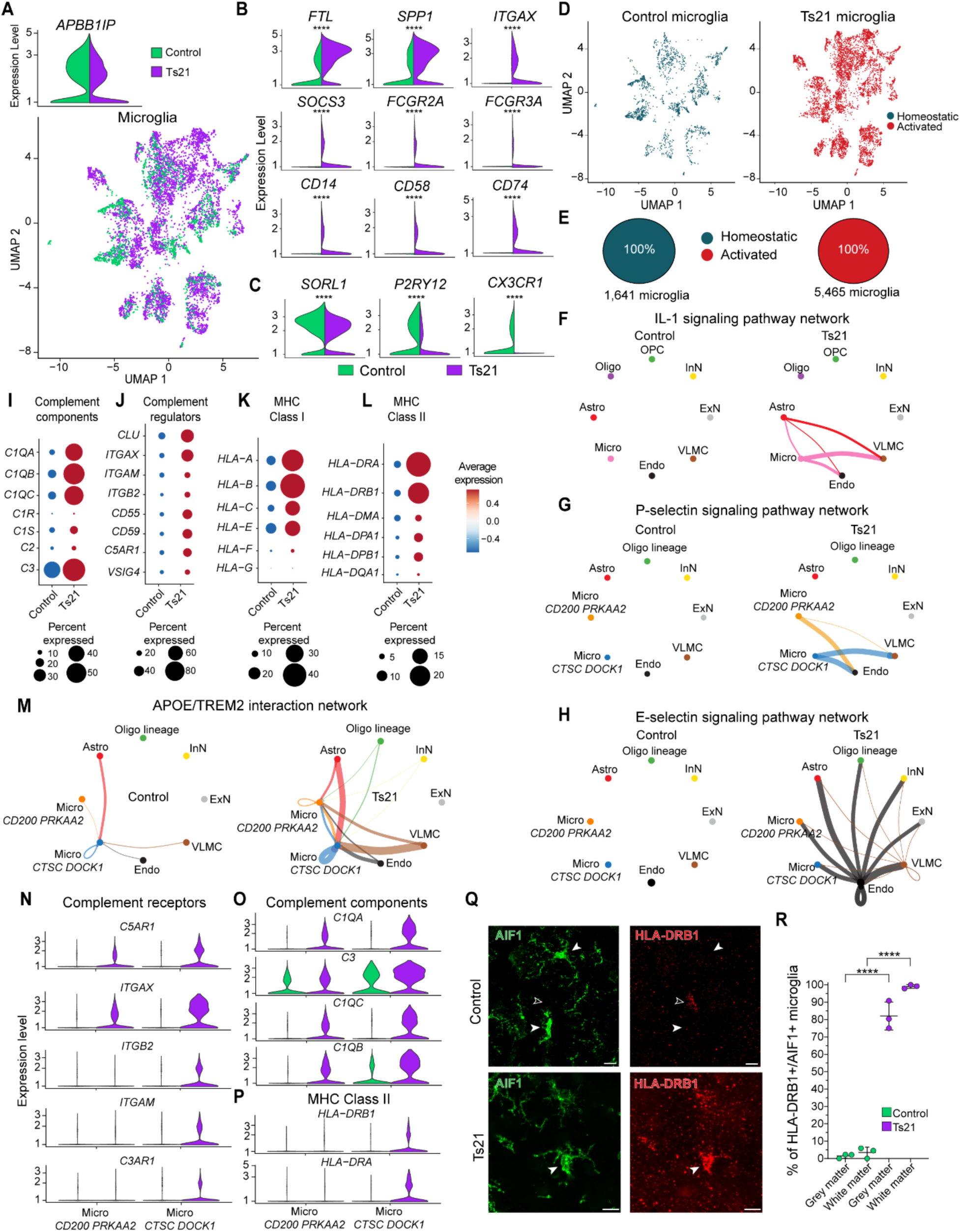
Dysregulated neuroimmune crosstalk and subtype-specific molecular features of Ts21 microglia (**A**) Upper panel: microglia were identified and subset by expression of the pan-astrocyte marker *APBB1IP*. Bottom panel: Transcriptomic UMAP visualization of all Control and Ts21 microglia analyzed in this study using a joint UMAP embedding, colored by condition. (**B and C**) Split gene expression violin plots of control (green) and Ts21 (purple) microglia for genes upregulated (B) or downregulated (C) in activated microglia. (**D**) Transcriptomic UMAP visualization of all Control (left) and Ts21 (right) microglia analyzed in this study using a joint UMAP embedding, colored by microglia state. (**E**) Proportion of control (left) and Ts21 (right) activated and homeostatic microglia. (**F to H**) Circle plot visualizing the inferred paracrine and autocrine signaling networks for IL-1 (F), P-selectin (G), and E-selectin (H). Each node represents a distinct cell subclass. Directed edges indicate statistically significant ligand–receptor-mediated interactions between cell groups where edge color corresponds to the sender (source) cell subclass and edge thickness reflects the relative interaction strength between cell groups. (**I to L**) Microglia gene expression dot plots of select genes from complement component (I), complement regulator (J), MHC class I (K), or MHC class II (L) HUGO gene families; dot size represents the percentage of nuclei expressing the gene and color represents the average expression value, (**M**) Circle plot visualizing the inferred APOE/TREM2 interaction network for control (left) and Ts21 (right). Directed edges indicate statistically significant ligand–receptor-mediated interactions between cell groups where edge color corresponds to the sender (source) cell subclass and edge thickness reflects the relative interaction strength between cell groups with equivalent maximum edge weight between control and Ts21. (**N to P**) Gene expression violin plots for complement receptors (N), complement components (O), and MHC class II (P). Violin plots are stratified by microglia subtypes: Micro *CD200 PRKAA2* and Micro *CTSC* DOCK1 for control microglia (green) and Ts21 microglia (purple). (**Q**) Representative images depicting AIF1+ positive microglia in the dorsolateral prefrontal cortex of control (upper panel) and Ts21 (lower panel) samples. The Ts21 panel highlights an AIF1+/HLA-DRB1+ microglial cell (white arrow), while the control panel shows AIF1+/HLA-DRB1-microglia (white arrows) and an AIF1-/HLA-DRB1+ intravascular immune cell (black arrow). (**R**) Quantification of the percentage of AIF1+ microglia expressing HLA-DRB1 in dlPFC grey matter or white matter of control (age: 2y, 4d postnatal) and Ts21 (age: 2y, 57d postnatal) samples. Each dot represents quantification from one tissue section (**** p<0.0001, two-tailed unpaired t-test). See also Supplementary Figs. 11 to 13 and Supplementary Table 5.

To investigate cell-autonomous effects underlying microglial activation, we employed multiomic gene regulatory network analysis to identify regulons underlying the activated state (**Supplementary Table 8**). This revealed increased activity of central inflammatory regulators, including PPARG, NFKB1, and the Chr21-encoded TF RUNX1 (**Supplementary Fig. 10**). Notably, there was decreased activity of the PPARG repressor regulon, indicating diminished suppression of inflammatory gene expression alongside the activation of pro-inflammatory pathways (**Supplementary Fig. 10**). Network analysis further revealed key TF-TG gene interactions associated with inflammation, including regulation of the Chr21-encoded interferon receptor *IFNGR2* by both RUNX1 and NFKB1 (**Supplementary Fig. 12A**). This finding suggests a Chr21-encoded positive feedback loop wherein RUNX1 directly regulates *IFNGR2* expression, exacerbating interferon signaling and inflammation.

Given the dysregulated, hyperactive interferon signaling observed across multiple Ts21 organ systems^62,63^, we next characterized the expression profiles of Chr21-encoded interferon receptors (*IL10RB*, *IFNGR2*, *IFNAR1*, *IFNAR2*, *IFNLR1*), non-Chr21-encoded interferon receptors (*IFNGR1*), and interferon response genes. This revealed that the vast majority of interferon receptors were highly upregulated across Ts21 cell types (**Supplementary Fig. 12B-C**). Furthermore, all cell classes demonstrated marked upregulation of interferon response genes, underscoring a broad and systemic interferon response within the early postnatal dlPFC (**Supplementary Fig. 12D**). Strikingly, vascular leptomeningeal cells (VLMCs) exhibited the highest number of upregulated interferon response genes, many of which were cell-type-specific, highlighting meningeal cells as a critical node for Ts21 inflammatory signaling (**Supplementary Fig. 12D**).

### Dysregulated neuroimmune crosstalk and subtype-specific molecular features of Ts21 microglia

Given the concurrent dysregulation of microglial and astrocytic cellular states, we systematically investigated cytokine-mediated signaling networks as non-cell-autonomous regulators of Ts21-associated neuroinflammation. Cell-cell interaction analyses identified significant enrichment of pro-inflammatory IL-1 and IL-6 signaling pathways, revealing inflammatory paracrine signaling between Ts21 astrocytes, microglia, and vascular-associated cells (**Fig. 7F and Supplementary Fig. 13A-B**). Notably, interaction profiling identified marked upregulation of Ts21 vascular pro-inflammatory selectin and galectin signaling (**Fig. 7G-H and Supplementary Fig. 13C-D**), which establish feedback loops through the stimulation and amplification of interleukin production^64–66^. E-selectin (*SELE*), an endothelial-specific adhesion molecule induced by cytokine activation^66^, was exclusively expressed in Ts21 vascular-associated cells (**Fig. 7G**). Furthermore, enhanced P-selectin signaling was identified in microglia-vascular interactions (**Fig. 7H**), reflecting its established role in immune cell modulation and the trafficking of inflammatory leukocytes along the vasculature^66^. Ts21 interaction networks reveal coordinated vascular-immune-glial cell crosstalk driving neuroinflammatory cascades through both soluble cytokine signaling and adhesion molecule-mediated cellular interactions, establishing a feedforward inflammatory axis between activated microglia and the cytokine-stimulated endothelium. Collectively, these findings identify critical regulators of microglial inflammation and interferon signaling while demonstrating a highly convergent interferon phenotype across cell types. These results also underscore the complex interplay between intrinsic Chr21-encoded factors—such as *RUNX1* and *IFNGR2*—and broader non-cell-autonomous inflammatory signaling as key drivers of Ts21 neuroinflammation in the early postnatal dlPFC.

Inflammatory signaling and microglial activation are particularly consequential during the early postnatal period, a critical window for synaptogenesis and complement-dependent synaptic pruning. Transcriptomic profiling revealed pronounced upregulation of complement cascade components and regulators—critical for synaptic tagging and engulfment—alongside elevated expression of MHC class I/II receptors in Ts21 microglia (**Fig. 7I-L**). These findings elucidate molecular signatures of enhanced complement-dependent synaptic tagging, engulfment, and phagocytic activity in Ts21 microglia, implicating dysregulated neuroimmune crosstalk in disrupted synaptic refinement during early brain development.

To further characterize microglial subtypes and identify populations with potential therapeutic relevance, we performed subclustering analysis which revealed two predominant microglia subtypes: Micro *CD200 PRKAA2* and Micro *CTSC DOCK1*, present in both control and Ts21 samples (**Supplementary Fig. 13E-I**). Importantly, both Ts21 subtypes exhibited increased interaction between the TREM2 receptor and APOE, a key pathway regulating the phagocytosis of apoptotic neurons and driving the transition of microglia from a homeostatic to an activated phenotype (**Fig. 7M**). Dysregulation of the TREM2/APOE axis has been implicated in exacerbating neuroinflammation, neuronal injury, and complement-mediated synaptic clearance, representing a critical therapeutic target in Alzheimer’s disease pathology^67^.

Transcriptomic profiling further revealed divergent molecular properties between microglial subtypes. Specifically, Ts21 Micro *CTSC DOCK1* demonstrated enriched expression of complement receptors critical for phagocytosis, including CR3 (ITGAM/ITGB2), CR5 (ITGAX/ITGB2), and C3aR (C3AR1), along with MHC class II components essential for antigen presentation (**Fig. 7N-P**). Notably, Ts21 Micro *CTSC DOCK1* exhibited increased expression of interferon receptors (**Supplementary Fig.12C**) and highly elevated expression of MHCII *HLA-DRB1*, one of the most consistently upregulated markers identified in Alzheimer’s disease^68^. Tissue immunohistochemical analyses confirmed significant increases in both the percentage of HLA-DRB1+/AIF1+ microglia and HLA-DRB1 immunointensity in the early postnatal Ts21 dlPFC, with more pronounced alterations observed in white matter (**Fig. 7Q-R, and Supplementary Fig. 13J-L**).

Together, these results demonstrate unique molecular signatures of activated microglial subtypes in Ts21, highlighting key functional differences in complement-mediated phagocytic activity and antigen presentation. These findings underscore the role of neuroimmune dysfunction in disrupted synaptic refinement during Ts21 early brain development and identify key pathways and cell subtypes for potential therapeutic intervention.

## Discussion

This study presents a comprehensive single-nucleus multiomic analysis of the early postnatal dorsolateral prefrontal cortex in Down syndrome, revealing a complex landscape of molecular and cellular dysregulation that underpins both the neurodevelopmental and the neurodegenerative features of this disorder. Our analyses identified broad and asymmetric increases in chromatin accessibility and gene expression across diverse cell types and chromosomes in Ts21. Notably, we uncovered shared patterns of differential accessibility, highlighting a pervasive epigenetic reorganization that extends beyond classical “gene dosage effects” and implicates both local and trans-acting regulatory mechanisms in molecular pathogenesis.

Our analysis also revealed multicellular dysfunction in the prefrontal cortex that converges at the level of the neurite and synapse. We identified molecular correlates of intrinsic pan-neuronal synaptic deficits, oligodendrocyte lineage progression and functional myelination defects, impaired neuron-glia adhesion, and activated antigen presenting microglia during a critical period of synaptogenesis (**Supplementary Fig. 14**). These findings provide a molecular framework for the circuit-level dysfunction observed in Down syndrome and highlight the vulnerability of neurite outgrowth, synaptic formation, and functional maturation during early postnatal brain development.

Accumulating evidence supports the classification of DS as a multisystem interferonopathy^62,63,69^. We identified a robust interferon response signature across all dlPFC cell types analyzed in this study, highlighting interferon activation at an early and critical stage of brain development (**Supplementary Fig. 12**). In addition, we identified a pronounced neuroinflammatory signature encompassing glial, immune, and vascular-associated cells along with cell-cell interaction networks and upstream transcriptional inflammatory mediators. Ts21 astrocytes exhibited a distinct shift toward a reactive phenotype, with upregulation of stress-responsive heat shock proteins and molecular chaperones, and a progressive increase in reactivity over the early postnatal period. We identified that this reactivity is driven in part by the activation of the key transcription factor AP-1 complex and is accompanied by the suppression of homeostatic gene expression and upregulation of cell-survival gene expression programs. Notably, the AP-1 complex has emerged as a potential therapeutic target bridging interferon signaling and downstream cellular pathology in Ts21 and represents a potential target for the modulation of astrocytes in Down syndrome^25^.

Furthermore, Ts21 microglia displayed marked activation in the early postnatal period, with increased expression of inflammatory markers, complement cascade components, and MHC class I/II receptors, as well as subtype-specific enrichment for phagocytic and antigen presentation pathways. We identified inflammatory mediators that drive this activation and further potentiate interferon receptor expression through regulation of the pro-inflammatory interferon receptor *IFNGR2*. This is especially relevant given interferon-responsive microglia have been shown to be highly phagocytic and actively shape cortical development^70^, suggesting that interferon-induced microglia pathologically and excessively prune synapses critical for physiological circuit maturation in the Ts21 dlPFC. Our analyses identify a Ts21 microglial subtype—Micro *CTSC DOCK1—*as an interferon-responsive microglial therapeutic target with potential to ameliorate synaptic and circuit dysfunction in DS. We further identified glial and immune state alterations to be amplified by dysregulated cytokine signaling and enhanced vascular-immune-glial crosstalk, establishing a multicellular feedforward inflammatory axis in the Ts21 brain.

DS is among the select few conditions that encompass both neurodevelopmental and neurodegenerative clinical features, with most individuals demonstrating progressive cognitive impairment by mid-adulthood^71,72^. While it is well-established that the neuropathological substrates of dementia precede clinical onset^73^, a central question is the timing of onset of these molecular changes. Strikingly, this study identified molecular signatures corresponding to six of eight established neurodegeneration hallmarks^74^—including metabolic dysfunction, oxidative stress, synaptic loss, neuroinflammation, impaired proteostasis, and activation of cell death pathways—during the early postnatal period in the Ts21 PFC. The concurrence of neurodevelopmental deficits and neurodegenerative signatures supports a model in which these processes are mechanistically interconnected, potentially compounding cognitive and functional impairment from the earliest stages of life in DS. These findings shift the temporal onset of DS-associated neurodegenerative processes to the early postnatal period and underscore the necessity for early biomarker development, therapeutic innovation, and clinical trials targeting disease modification within this critical developmental window.

## Acknowledgements

We thank H. Thurston, A. Osterman, M. Jandy, N. West, and M. Russo for technical assistance and critical discussion of the data; K. Knobel at the Waisman IDD Model Core for core services. This work was supported by the National Institutes of Health grants 1R01HD106197 (to A.M.M.S., D.W., S-C. Z., and A.B.), R01AG067025 and RF1MH128695 (to D.W.), as well as 1F30MH140382-01 and the Medical Scientist Training Program grant T32 GM140935 (to R.D.R), and P50HD105353 (to the Waisman Center). Funding was also provided by the Brain Research Foundation BRFSG-2023-11 (to A.M.M.S.), and the Morse Society Fellowship to R.D.R.

## Authors contributions

R.D.R., S-C. Z., A.B., and A.M.M.S. conceived and designed the study. R.D.R., M.H., R.J.C., D.K.S., C.W., E.G., and A.M.M.S. performed the experiments. K.H.A., S.K., P.K., X.H., J.S., and D.W. analyzed the genomics data. R.D., K.H.A., S.K., M.H., and A.M.M.S. wrote the manuscript. All authors edited the manuscript.

## Data and materials availability

The genomic data has been submitted to dbGaP (accession number pending). The data can be interactively visualized at https://daifengwanglab.shinyapps.io/DS_PFC/. All other data are available in the main paper or supplementary information.

---

**Supplementary Fig. 1.**
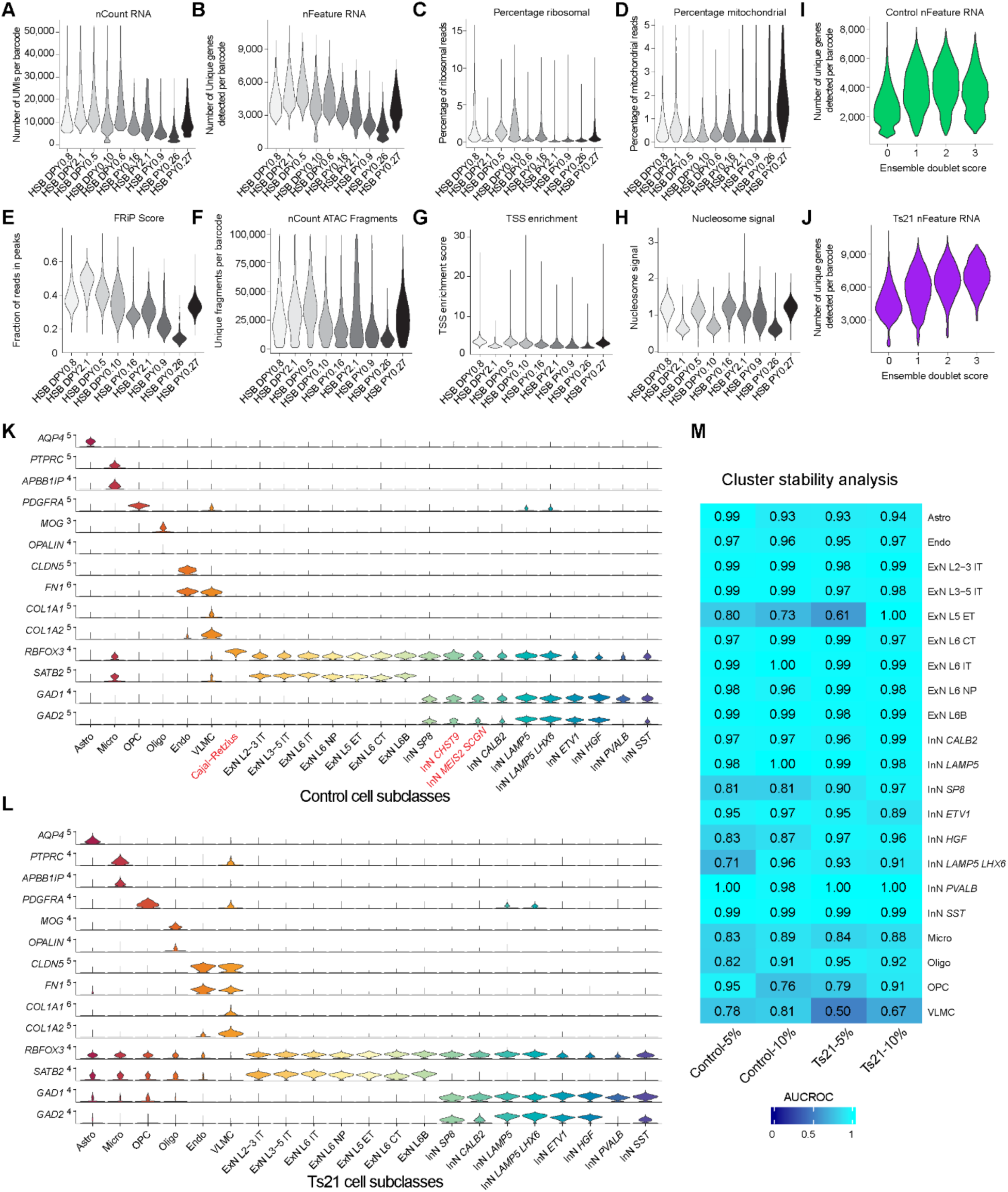
Single-nucleus multiomic gene expression and chromatin accessibility quality control and cell-type annotation. (**A to D**) Gene expression quality control metrics following snRNA-seq data processing (Methods). Each violin plot represents one biological sample. (**E to H**) ATAC-seq quality control metrics following snATAC-seq data processing (Methods). Each violin plot represents one biological sample. (**I and J**) Violin plots depicting the number of unique genes detected per barcode for control (I) and Ts21 (J) ensemble doublet scores. Nuclei were classified as doublets and removed if they were flagged by at least two of the three doublet detection methods (Methods). (**K and L**) Violin plots of cell type annotation marker genes for control (K) and Ts21 (L) cell subclasses. (**M**) Cluster stability analysis of control and Ts21 cell subclasses with neighbor voting-based approach. Individual cell subclasses were subset with 5% or 10% of the original sample and area under the receiver operating characteristic (AUROC) scores for each cell subclass depicted in blue.

**Supplementary Fig. 2.**
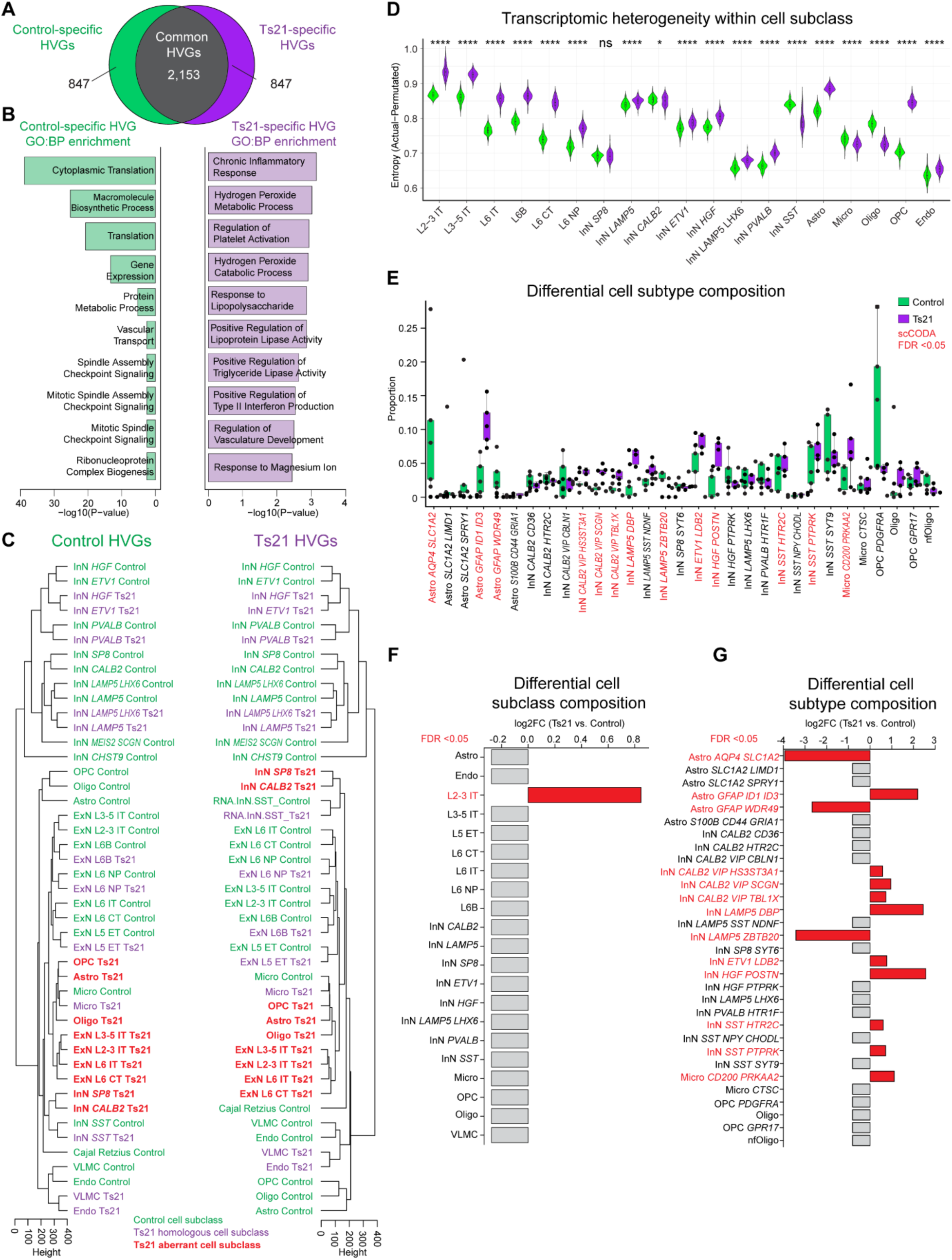
Condition-specific highly variable genes, transcriptomic heterogeneity, and differential cell composition analysis. (**A**) Venn diagram depicting the number of overlapping highly variable genes (HVGs). The top 3,000 highly variable genes were calculated for control and Ts21 independently and the number of shared and condition-specific HVGs compared. (**B**) The top 10 Gene Ontology: biological process enrichment terms for control-specific (left) or Ts21-specific (right) HVGs, ranked by P-value. (**C**) Dendrograms depicting unsupervised hierarchical clustering of pseudobulked nuclei, stratified by cell subclass and condition. This analysis was conducted independently using the top 3,000 HVGs calculated for control (left) or Ts21 (right). (**D**) Violin plots depicting transcriptomic heterogeneity, defined as unstructured entropy subtracted from the actual entropy, for control (green) and Ts21 (purple) cell subclasses (* p<0.05, **** p<0.0001, unpaired Wilcoxon rank sum test). (**E**) Differential cell subtype composition analysis with age and sex as covariates at the cell subtype annotation level. Statistically significant compositional changes for individual cell subtypes are highlighted in red (FDR <0.05). (**F and G**) Bar plots depicting cell compositional changes at the cell subclass (F) and cell subtype (G) level. Each bar represents the log2 fold change, with positive values indicating increased abundance in Ts21 and negative values indicating decreased abundance in Ts21. Red bars represent statistically significant changes (FDR <0.05) and grey bars represent non-significance.

**Supplementary Fig. 3.**
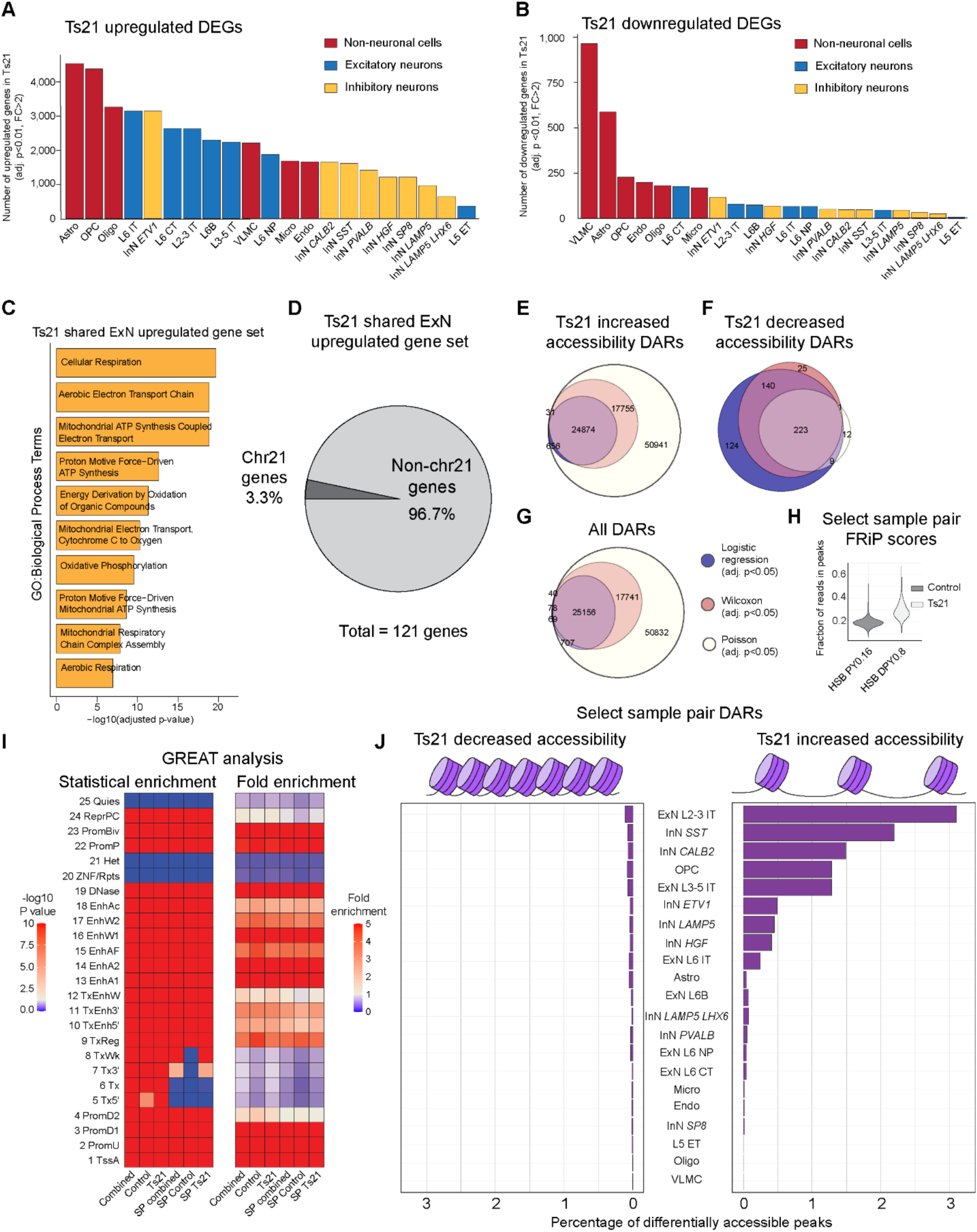
Differential gene expression and differentially accessible region (DAR) analyses. (**A and B**). Bar plots depicting the number of highly upregulated (A, adjusted p-value <0.01, FC>2) or highly downregulated (B, adjusted p-value <0.01, FC>2) genes in Ts21, stratified by cell subclass and bars colored by major cell class. (**C**) The top 10 Gene Ontology: biological process enrichment terms for Ts21 highly upregulated genes shared across all Ts21 cell subclasses, ranked by adjusted p-value. (**D**) Pie chart depicting the percentage of Ts21 highly upregulated genes shared across all cell subclasses that are encoded on Chr21. (**E to G**) Venn diagrams depicting the number of differentially accessible regions (DARs) calculated with three independent peak-calling methods: logistic regression, Wilcoxon rank sum, and Poisson generalized linear model (adjusted p-value <0.05, all methods). The most conservative method, logistic regression, was utilized for downstream analysis. (**H)** Fraction of reads in peaks (FriP) scores for additional select matched-sample analyses at the earliest study timepoint (HSBPY0.16, control 60 days postnatal; HSBDPY0.8, Ts21 65 days postnatal). (**I**) Heatmaps depicting results from the peak-set Genomic Regions Enrichment of Annotations Tool (GREAT) analysis. Statistical enrichment (left heatmap, p-value calculated using a negative binomial distribution) and fold enrichment (right heatmap) were determined for each state relative to the total peak set. “Combined”, combined whole dataset peak set; “SP”, selected sample pair peak set. (**J**) Bar plot of the selected sample pair percentage of differentially accessible regions (DARs) per cell subclass with an adjusted p-value threshold of 0.05 from logistic regression. The percentage of DARs is defined as the number of DARs relative to the total number of peaks with observed fragments in at least 1% of barcodes for each cell type.

**Supplementary Fig. 4.**
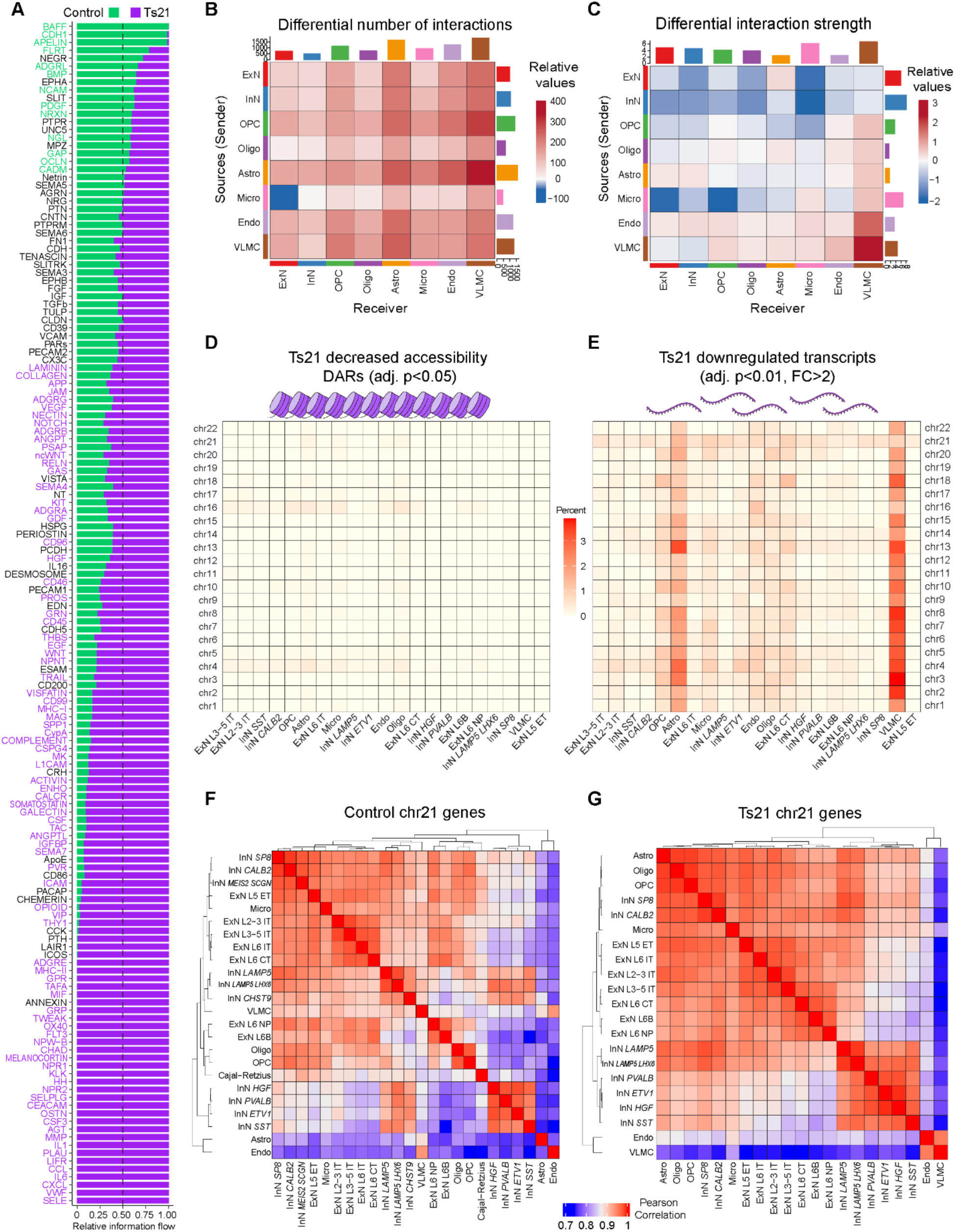
Cell-cell interactions networks and chromosomal patterns of Ts21 differential chromatin accessibility and gene expression. (**A**) Relative information flow plots depicting the aggregate network weights of all cell-cell interactions identified in this study. Interaction pathways are arranged vertically according to differential communication strength, with horizontal bars representing normalized information flow values. Pathways significantly enriched in control are shown in green, while those significantly enriched in Ts21 are indicated in purple (p<0.05, paired Wilcoxon test). (**B**) Heatmap illustrating the differential number of interactions among distinct cell types, Ts21 vs. control. The y-axis depicts the source (sender) cells, and the x-axis depicts the receiver cells. The top bar plot displays the sum of absolute values for each column, representing the total change in incoming signaling to each cell population. The right bar plot shows the sum of absolute values for each row, indicating the total change in outgoing signaling. In the heatmap color scale, red denotes a relative increase in the number of interactions in Ts21. (**C**) Heatmap illustrating the differential strength of interactions (differential edge weight) among distinct cell types, Ts21 vs. control. The y-axis depicts the source (sender) cells, and the x-axis depicts the receiver cells. The top bar plot displays the sum of absolute values for each column, representing the total change in incoming signaling strength to each cell population. The right bar plot shows the sum of absolute values for each row, indicating the total change in outgoing signaling strength. In the heatmap color scale, red denotes a relative increase in interaction strength in Ts21. (**D**) Heatmap of the percentage of differentially accessible regions (DARs) with decreased accessibility in Ts21 plotted by cell subclass and autosome number (adjusted p<0.05, logistic regression). The numerator is the number of DARs, and the denominator is the number of combined peaks where a 1% fraction of barcodes showed a >0 number of fragments aligned. (**E**) Heatmap of the percentage of highly downregulated genes in Ts21 plotted by cell subclass and autosome number (adjusted p<0.01, log2FC <-1, NEBULA). (**F and G**) Heatmap depicting pairwise Pearson correlations of Chr21 gene expression on subclass-level pseudobulked nuclei for control (F) and Ts21 (G).

**Supplementary Fig. 5.**
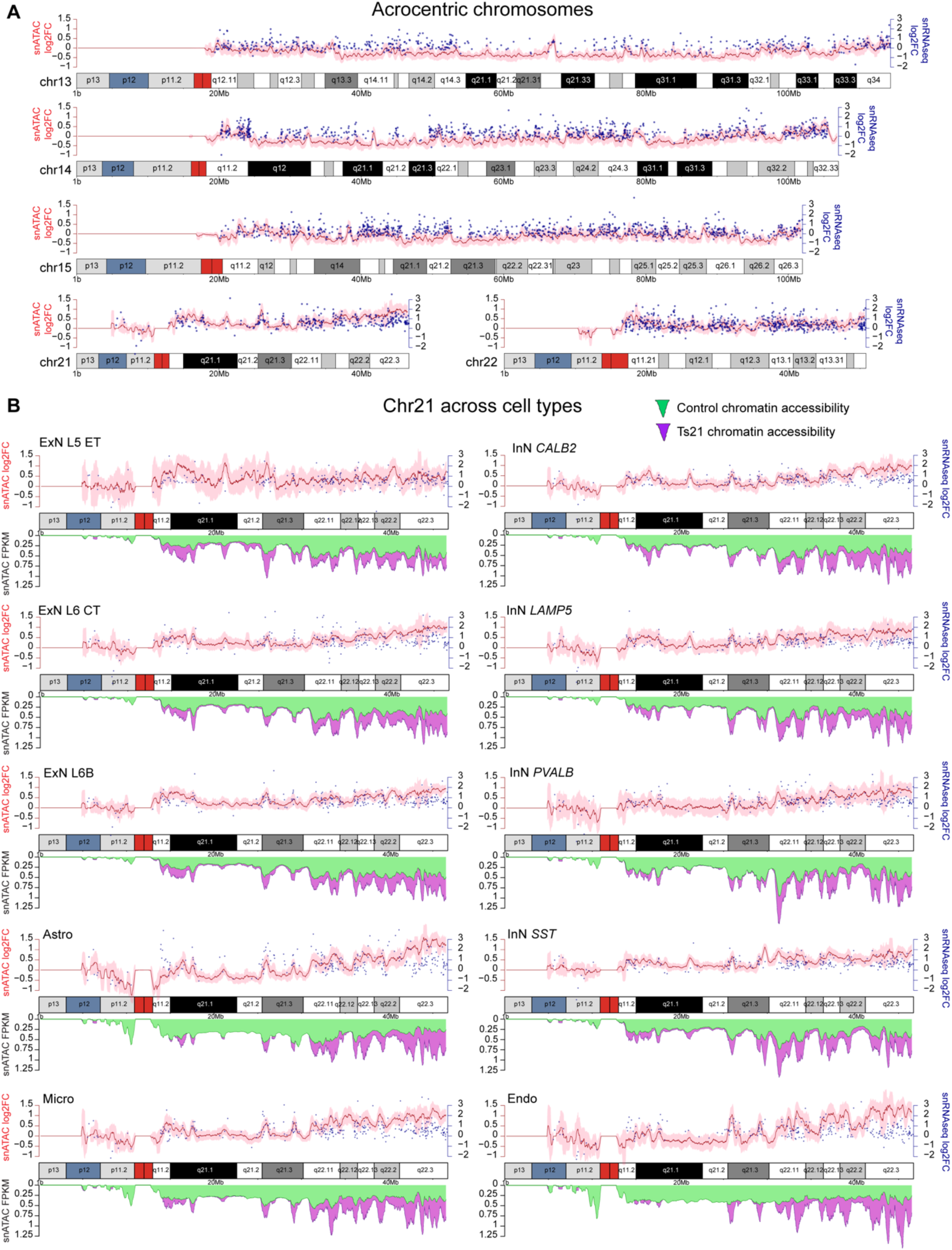
Chromosome-scale multiomic gene expression and chromatin accessibility across cell subtypes. (**A**) Integrated visualization of acrocentric chromosome gene expression and chromatin accessibility at the megabase pair (Mbp) scale for intratelencephalic excitatory neurons (ExN L2-3 IT, ExN L3-5 IT, ExN L6 IT) in pseudobulk. Below, ideograms for human acrocentric chromosomes depicting chromosome length (Mbp) and cytoband patterns. Above, the log fold change of accessibility (in FPKM) for uniform 10 kbp bins with a rolling mean (red line) and standard deviation across 40 bins (pink shading). Mean ExN IT differential gene expression log fold change for individual genes are depicted at gene transcription start site (TSS) coordinates (blue dots). (**B**) Integrated visualization of Chr21 gene expression and chromatin accessibility for select cell subclasses. Center, Chr21 ideogram depicting chromosome length and cytoband patterns. Below, snATAC FPKM coverage for control (green) and Ts21 (purple). Above, the log fold change of accessibility (in FPKM) for uniform 10 kbp bins with a rolling mean (red line) and standard deviation across 40 bins (pink shading). Mean cell subclass differential gene expression log fold change for individual genes are depicted at gene transcription start site (TSS) coordinates (blue dots).

**Supplementary Fig. 6.**
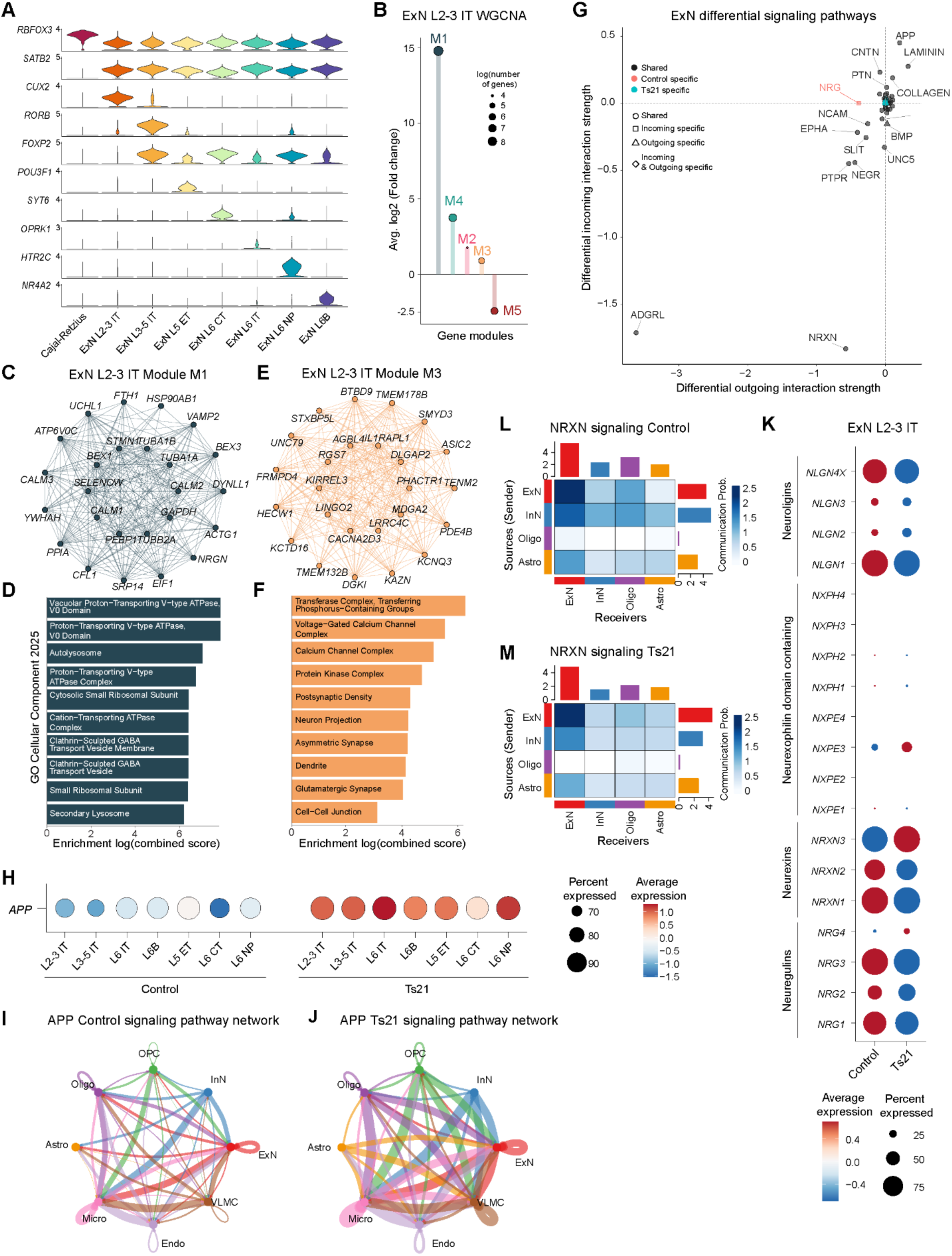
Excitatory neuron gene co-expression and cell-cell interaction networks. (**A**) Violin plots depicting marker gene expression of excitatory neuron subtypes. ExN, excitatory neuron; IT, intratelencephalic; ET, extratelencephalic; CT, corticothalamic; NP, near-projecting. (**B**) Bar plot depicting differential gene co-expression module enrichment between Ts21 and control ExN L2-3 IT. Positive y-axis values indicate module fold enrichment in Ts21 and circle size depicts the log number of genes within each module. (**C**) Co-expression network visualization of Module 1 for ExN L2-3 IT neurons with the top 25 module hub genes visualized. (**D**) Bar plot depicting gene set enrichment results for ExN L2-3 IT Module 1 co-expression network with the top Gene Ontology: Cellular Components (GO:CC) enrichment terms shown. (**E**) Co-expression network visualization of Module 3 for ExN L2-3 IT neurons with the top 25 module hub genes visualized. (**F**) Bar plot depicting gene set enrichment results for ExN L2-3 IT Module 4 co-expression network with the top Gene Ontology: Cellular Components (GO:CC) enrichment terms shown. (**G**) Scatter plot depicting differential outgoing and incoming signaling changes between Ts21 and control for all ExN in pseudobulk. Each point represents a signaling pathway, with its position along the x-axis indicating the change in outgoing interaction strength and the y-axis indicating the change in incoming interaction strength. Positive values denote increased signaling in Ts21 relative to control. (**H**) *APP* expression dot plots for ExN subclasses. (**I and J**) Circle plots visualizing the inferred APP interaction network for control (I) and Ts21 (J). Directed edges indicate statistically significant ligand–receptor-mediated interactions between cell groups where edge color corresponds to the sender (source) cell subclass and edge thickness reflects the relative interaction strength between cell groups with equivalent maximum edge weight between control and Ts21. (**K**) Dot plots of neurexin, neuregulin, neuroligins, and neurexophilin domain containing gene family expression in control (left) and Ts21 (right) ExN L2-3 IT. (**L and M**) Heatmaps of control (L) and Ts21 (M) NRXN interaction strength networks between source and target cell groups, where color intensity denotes interaction magnitude. Bar plots summarize the total outgoing (above) and incoming (right) NRXN interaction strengths for each cell group.

**Supplementary Fig. 7.**
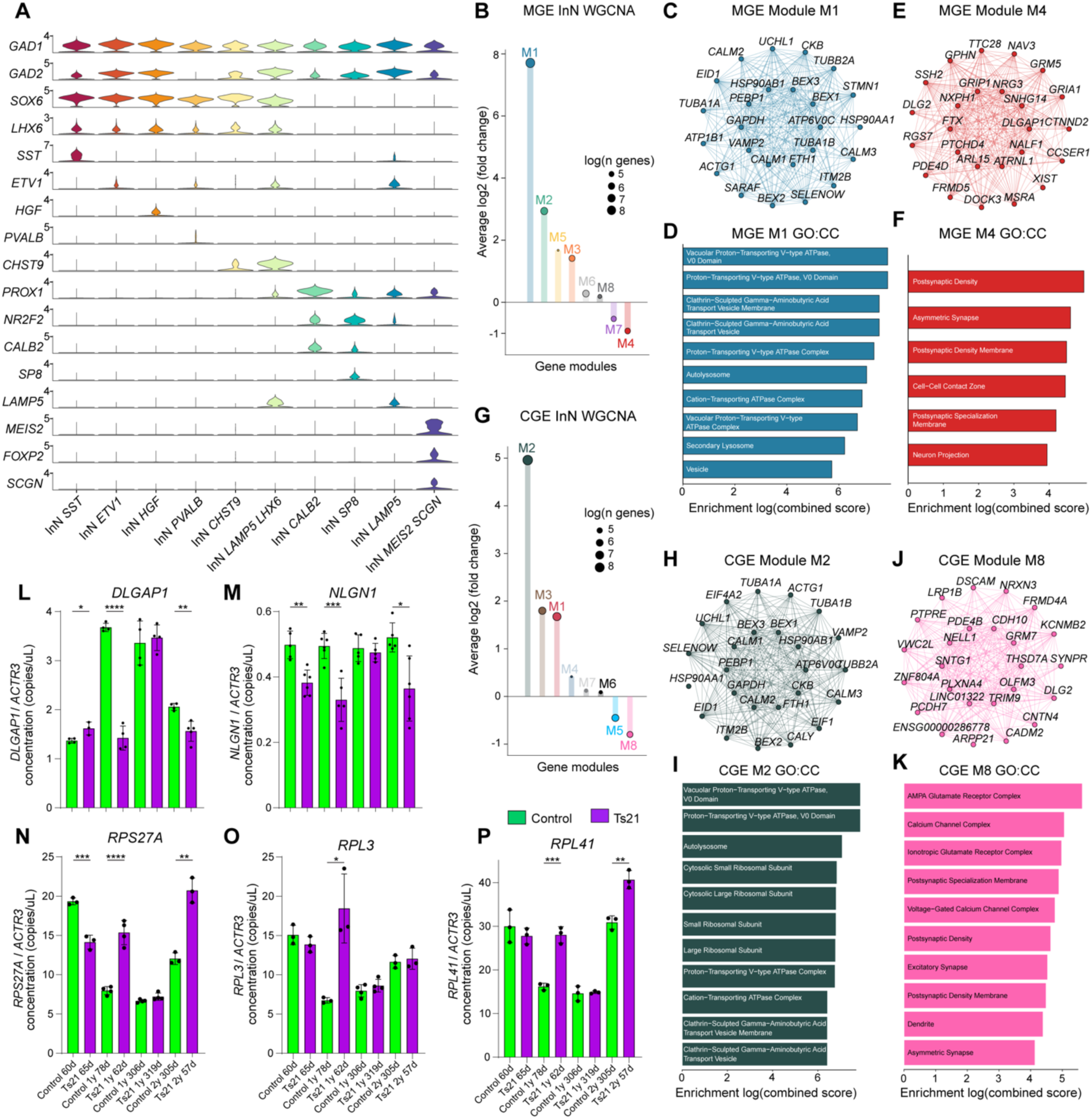
Inhibitory neuron gene co-expression networks and ddPCR validations. (**A**) Violin plots depicting marker gene expression of inhibitory neuron subtypes. (**B**) Bar plot depicting differential gene co-expression module enrichment between Ts21 and control MGE InN. Positive y-axis values indicate module fold enrichment in Ts21 and circle size depicts the log number of genes within each module. (**C**) Co-expression network visualization of medial ganglionic eminence (MGE)-derived neuron Module 1 with the top 25 module hub genes visualized. (**D**) Bar plot depicting gene set enrichment results for MGE Module 1 co-expression network with the top Gene Ontology: Cellular Components (GO:CC) enrichment terms shown. (**E**) Co-expression network visualization of MGE Module 4 with the top 25 module hub genes visualized. (**F**) Bar plot depicting gene set enrichment results for MGE Module 4 co-expression network with the top Gene Ontology: Cellular Components (GO:CC) enrichment terms shown. (**G**) Bar plot depicting differential gene co-expression module enrichment between Ts21 and control CGE InN. Positive y-axis values indicate module fold enrichment in Ts21 and circle size depicts the log number of genes within each module (**H**) Co-expression network visualization of caudal ganglionic eminence (CGE)-derived neuron Module 2 with the top 25 module hub genes visualized. (**I**) Bar plot depicting gene set enrichment results for CGE Module 2 co-expression network with the top Gene Ontology: Cellular Components (GO:CC) enrichment terms shown. (**J**) Co-expression network visualization of CGE Module 8 with the top 25 module hub genes visualized. (**K**) Bar plot depicting gene set enrichment results for CGE Module 8 co-expression network with the top Gene Ontology: Cellular Components (GO:CC) enrichment terms shown. (**L to P**) Bar plots depicting ddPCR-quantified expression of synaptic (L and M) and ribosomal (N to P) genes from control (green) and Ts21 (purple). Concentration of target genes was normalized to housekeeping control *ACTR3* (* p<0.05, ** p<0.01, *** p<0.001, **** p<0.0001, two-tailed unpaired t-test).

**Supplementary Fig. 8.**
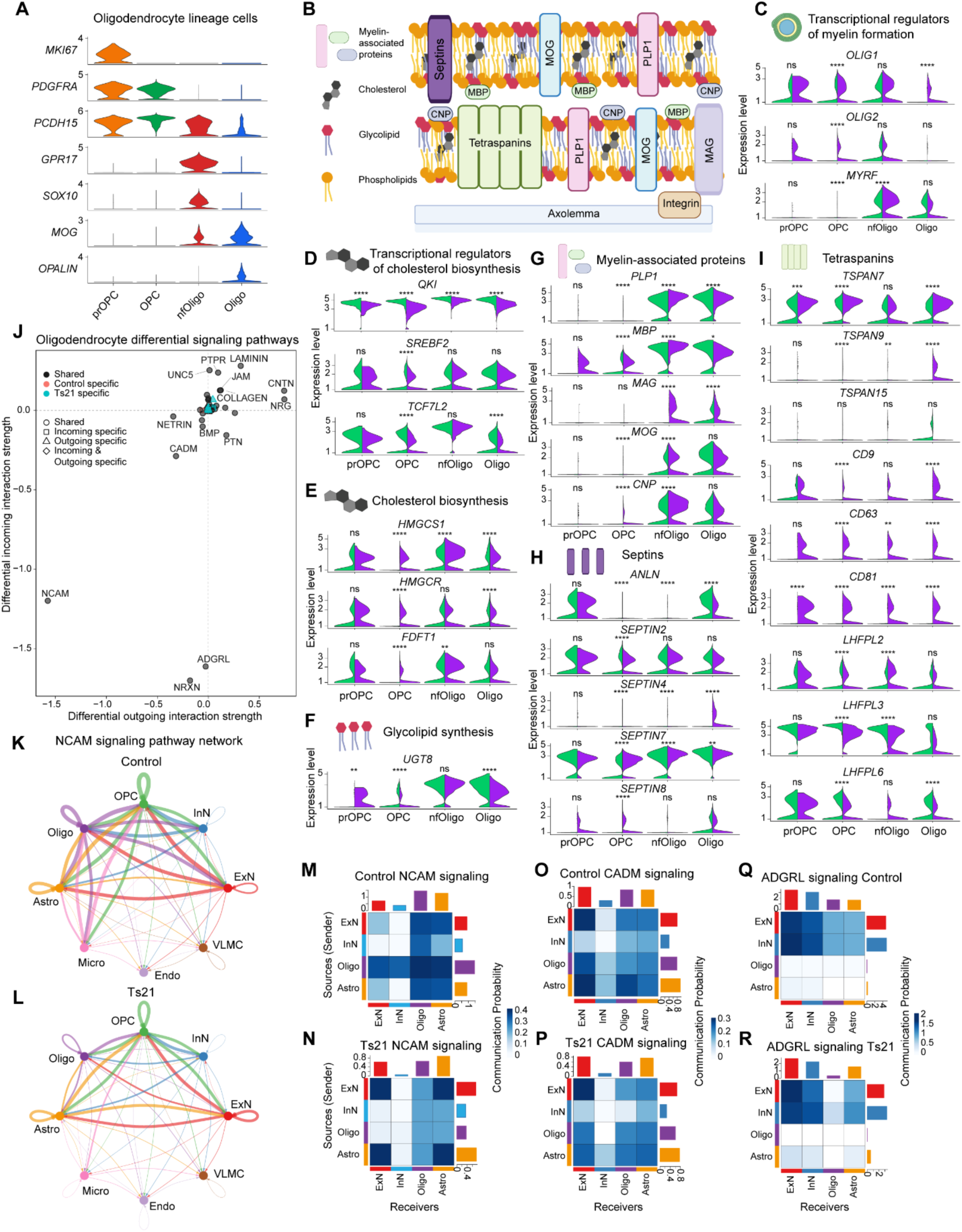
Oligodendrocyte cell-cell interaction networks and expression of myelin-related genes. **(A)** Violin plots depicting marker gene expression of oligodendrocyte lineage cells. prOPC, proliferative oligodendrocyte precursor cells; OPC, oligodendrocyte precursor cells; nfOligo, newly formed oligodendrocyte; Oligo, oligodendrocyte. (**B**) Schematic of major myelin components (top) and axolemma (bottom). (**C to I**) Split violin plots of genes encoding select transcriptional myelin regulators (C), transcriptional cholesterol regulators (D), cholesterol biosynthesis components (E), glycolipid synthesis components (F), myelin-associated protein (G), septins (H), and tetraspanins (I) plotted by oligodendrocyte lineage cell subclass. Control and Ts21 gene expression violin plots are green and purple, respectively. (* p<0.05, ** p<0.01, *** p<0.001, **** p<0.0001, Wilcoxon Rank Sum test). (**J**) Scatter plot depicting differential outgoing and incoming signaling changes between Ts21 and control for all oligodendrocytes (including nfOligo and Oligo) in pseudobulk. Each point represents a signaling pathway, with its position along the x-axis indicating the change in outgoing interaction strength and the y-axis indicating the change in incoming interaction strength. Positive values denote increased signaling in Ts21 relative to control. (**K and L**) Circle plots visualizing the inferred NCAM interaction network for control (K) and Ts21 (L). Directed edges indicate statistically significant ligand–receptor-mediated interactions between cell groups where edge color corresponds to the sender (source) cell subclass and edge thickness reflects the relative interaction strength between cell groups with equivalent maximum edge weight between control and Ts21. (**M to R**) Heatmaps of control (top panels) and Ts21 (bottom panels) NCAM (M and N), CADM (O and P), and ADGRL (Q and R) interaction strength networks between source and target cell groups. Color intensity denotes interaction magnitude and bar plots summarize the total outgoing (above) and incoming (right) interaction strengths for each cell group.

**Supplementary Fig. 9.**
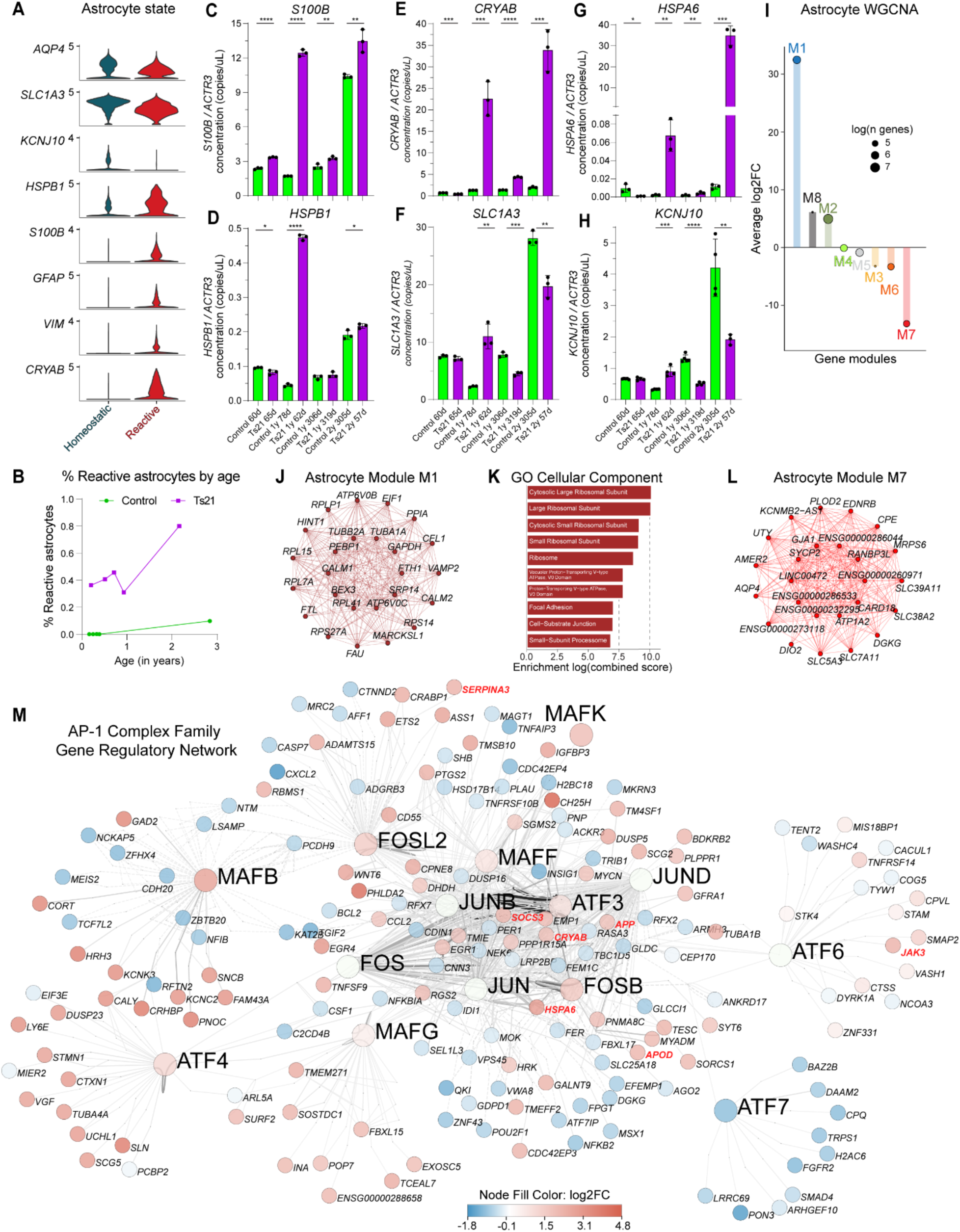
Reactive astrocyte ddPCR, astrocyte gene co-expression networks, and astrocyte AP-1 complex gene regulatory network. **(A)** Violin plots depicting marker gene expression of astrocyte state annotations. Pan-astrocyte marker *AQP4*. Reactive astrocytes were annotated based on increased *HSPB1*, *GFAP*, *VIM*, and *CRYAB* expression with concomitant decreased expression of *SLC1A3* and *KCNJ10*. (**B**) Percentage of astrocytes that were identified as reactive plotted by age of control (green) and Ts21 (purple) samples. (**C to H**) Bar plots depicting ddPCR-based validation of genes with increased expression in reactive astrocytes (C to E, and G) or decreased expression in reactive astrocytes (F and H) from control (green) and Ts21 (purple) samples. Concentration of target genes was normalized to housekeeping control *ACTR3* (* p<0.05, ** p<0.01, *** p<0.001, **** p<0.0001, two-tailed unpaired t-test). (**I**) Bar plot depicting differential gene co-expression module enrichment between Ts21 and control astrocytes. Positive y-axis values indicate module fold enrichment in Ts21 and circle size depicts the log number of genes within each module. (**J**) Co-expression network visualization of astrocyte Module 1 with the top 25 module hub genes visualized. (**K**) Bar plot depicting gene set enrichment results for astrocyte Module 1 co-expression network with the top Gene Ontology: Cellular Components (GO:CC) enrichment terms shown. (**L**) Co-expression network visualization of astrocyte Module 7 with the top 25 module hub genes visualized. (**M**) AP-1 complex family gene regulatory network in astrocytes. The network depicts the connections between AP-1 complex family eRegulons, their associated epigenomic regions, and target genes. For each AP-1 complex family eRegulon, the top 10 upregulated and top 10 downregulated target genes were identified based on astrocyte differential gene expression. Each node represents a gene and is colored according to differential expression log2FC in Ts21 astrocytes. Node label size indicates type: larger text denotes AP-1 eRegulons, while smaller italicized text denotes target genes. Solid edges represent activating interactions, and dashed edges indicate repressive interactions. Reactive astrocyte and neuroinflammation-associated target genes are highlighted in red.

**Supplementary Fig. 10.**
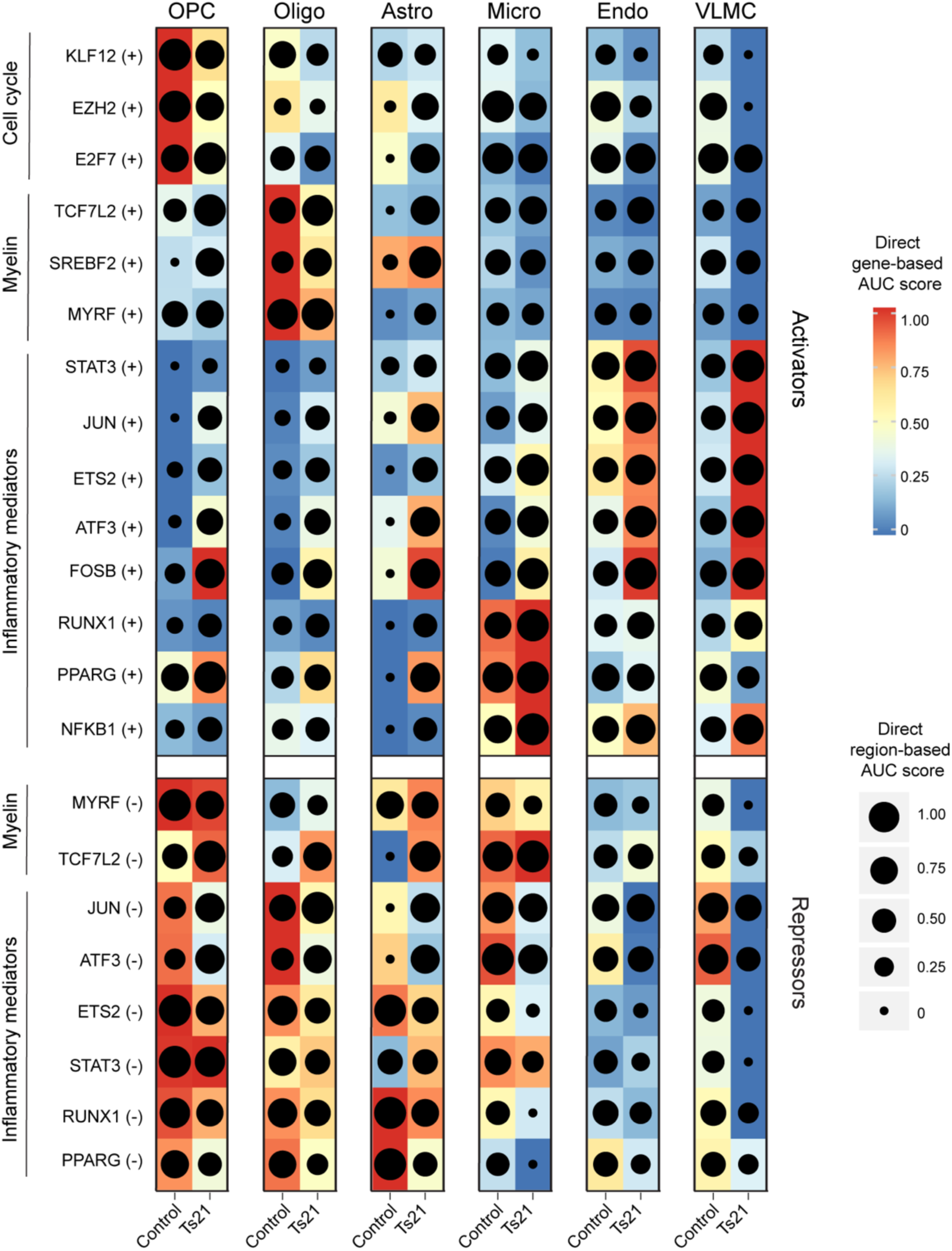
Multiomic gene regulatory network of Control and Ts21 non-neuronal cells Combined heatmap and dot plot depicting select SCENIC+ enhancer-driven regulon (eRegulon) activities from the non-neuronal cell SCENIC+ gene regulatory network. Heatmap colors indicate the gene expression-based AUC scores and dot sizes are scaled based on epigenomic region-based AUC scores. The eRegulons are split based on whether they are transcriptional activators (top, + regulons) or repressors (bottom, - regulons) and grouped based on broad functional annotations (cell cycle-related, myelin-related, and inflammatory mediator regulons).

**Supplementary Fig. 11.**
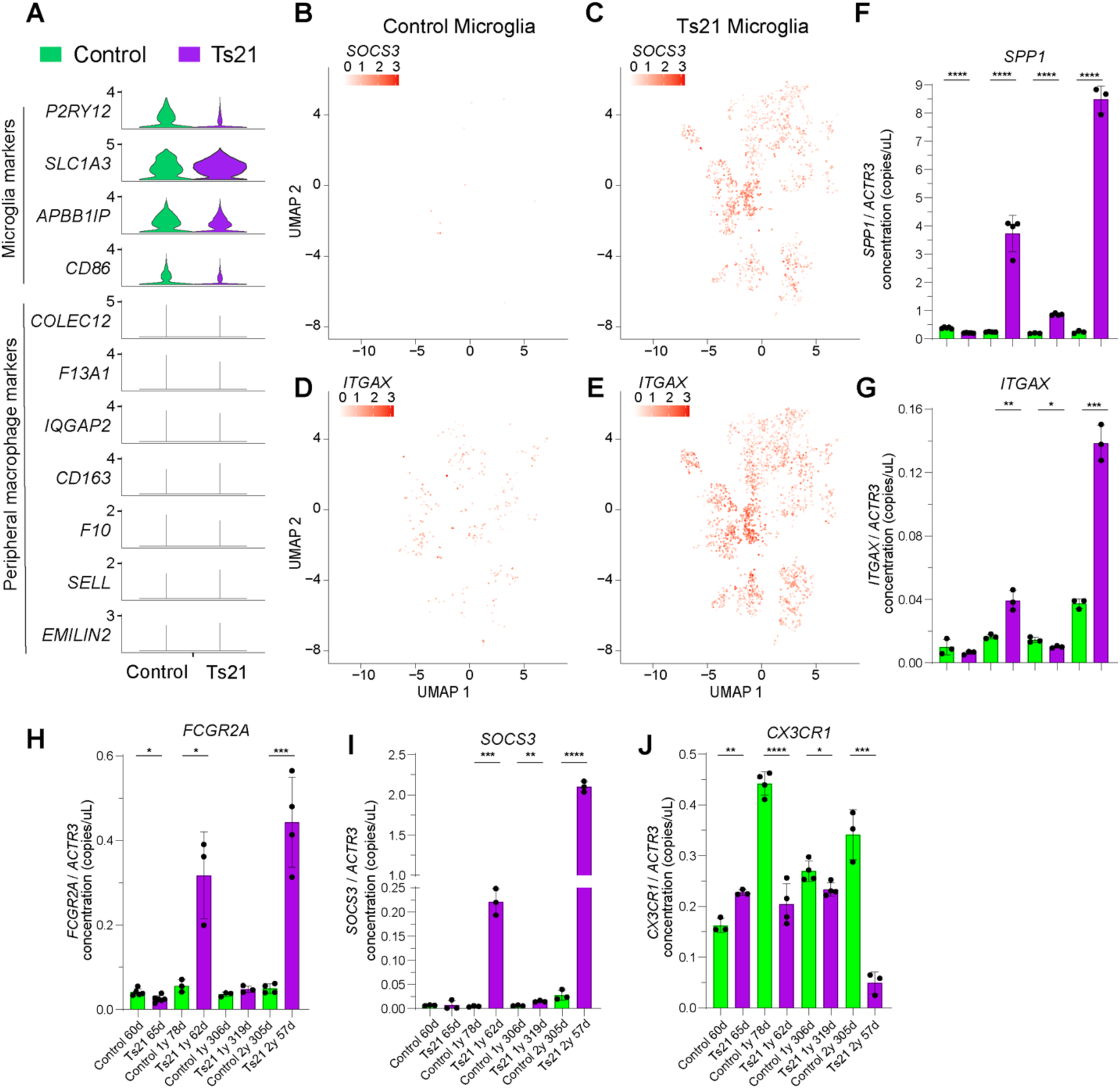
Gene expression profiles of Ts21 microglia **(A)** Violin plots depicting gene expression of microglia markers (top) or peripheral macrophage markers (bottom) for control (left) and Ts21 (right). (**B to E**) UMAP feature plot visualizing expression levels of the activated microglia markers *SOCS3* (B and C) and *ITGAX* (D and E) in control (left) and Ts21 (right microglia). Each point represents a single nuclei position on the microglia-only UMAP, with color intensity reflecting normalized gene expression. (**F to J**) Bar plots depicting ddPCR-based validation of genes with increased expression in activated microglia (F to I) or decreased expression in activated microglia (J) from control (green) and Ts21 (purple) samples. Concentration of target genes was normalized to housekeeping gene *ACTR3* (* p<0.05, ** p<0.01, *** p<0.001, **** p<0.0001, two-tailed unpaired t-test).

**Supplementary Fig. 12.**
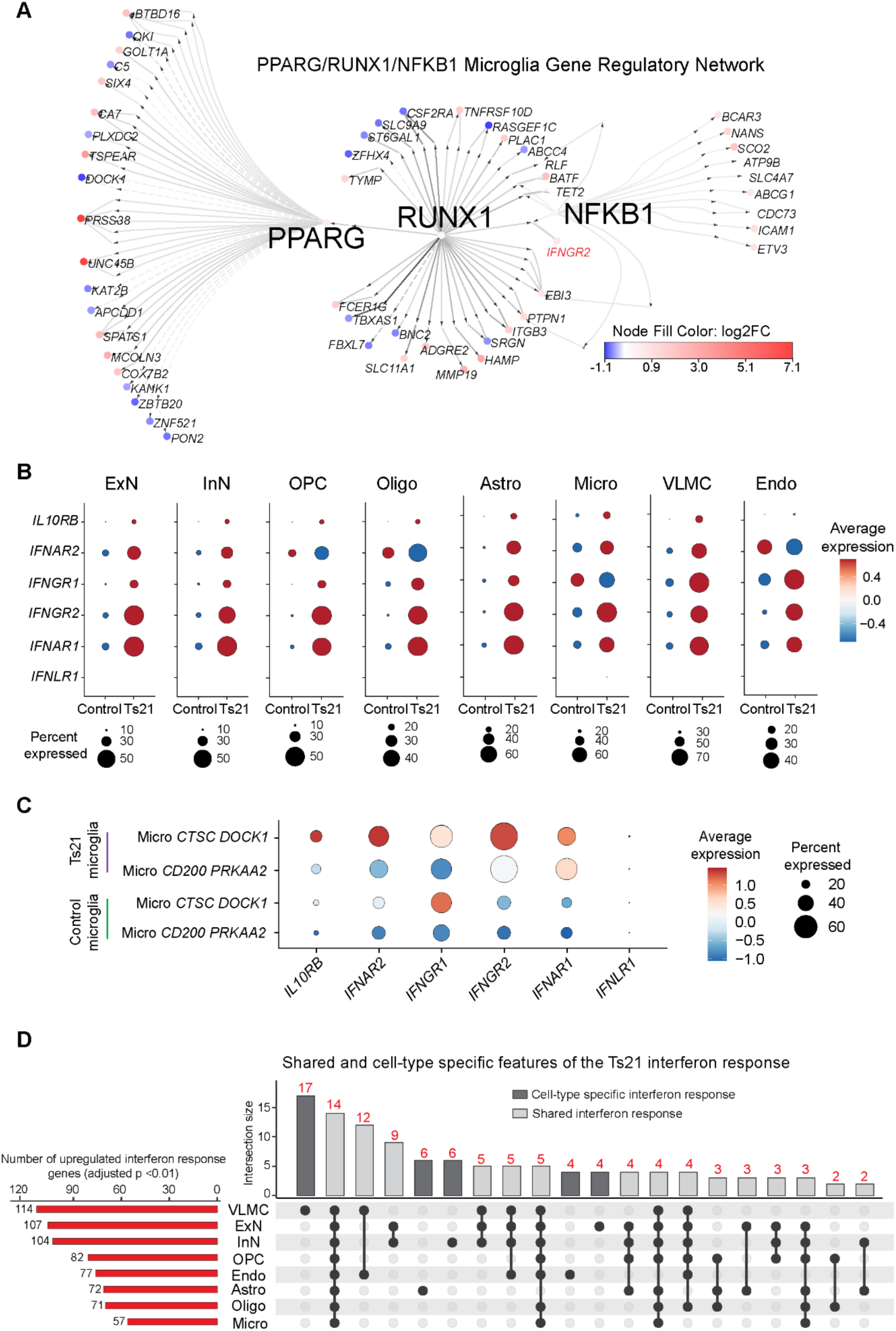
Microglia neuroinflammatory-associated gene regulatory network and cell-type-specific features of the Ts21 interferon response. (**A**) PPARG, RUNX1, NFKB1 regulatory network visualization for Microglia. The network depicts the connections between inflammatory mediators PPARG, RUNX1, and NFKB1 eRegulons, their associated epigenomic regions, and target genes. For each eRegulon, the top 10 upregulated and top 10 downregulated target genes were identified based on microglia differential gene expression. Each node represents a gene and is colored according to differential expression log2FC in Ts21 microglia. Node label size indicates type: larger text denotes eRegulons, while smaller italicized text denotes target genes. Solid edges represent activating interactions, and dashed edges indicate repressive interactions. The regulation of *IFNGR2* by RUNX1 is highlighted in red. (**B**) Dot plots visualizing interferon receptor gene expression across conditions and cell types. (**C**) Dot plots visualizing interferon receptor gene expression across Ts21 (top) and control (bottom) microglial subtypes (**D**) Upset plot of upregulated interferon response genes per cell type (adjusted p<0.01, NEBULA). The bottom panel depicts a matrix layout where filled dots indicate the sets included in each intersection, connected by lines to highlight the combination. The top bar chart quantifies the number of upregulated interferon response genes in each intersection. The left bar chart depicts the set size for each cell type. Bars are shaded on the cell-type specificity of the differentially expressed interferon response gene set.

**Supplementary Fig. 13.**
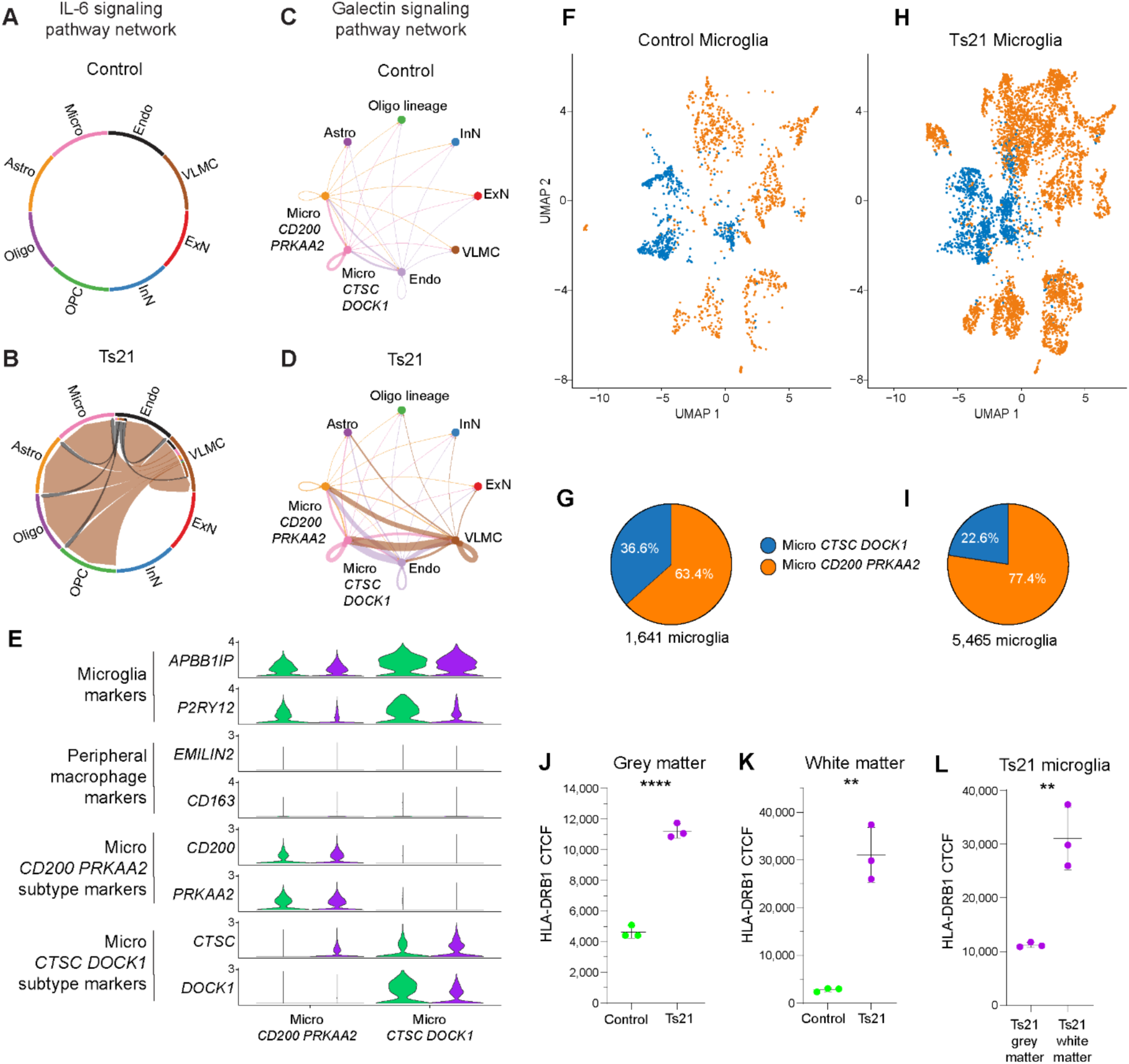
Ts21 vascular-glial-immune signaling and microglial subtype-specific features. (**A and B**) Chord diagrams visualizing the inferred IL-6 interaction network for control (A) and Ts21 (B). Chords, representing directed edges, indicate statistically significant ligand–receptor-mediated interactions between cell groups where chord color corresponds to the sender (source) cell subclass and chord thickness reflects the relative interaction strength between cell groups with equivalent maximum edge weight between control and Ts21. (**C and D**) Circle plots visualizing the inferred galectin interaction network for control (C) and Ts21 (D). Directed edges indicate statistically significant ligand–receptor-mediated interactions between cell groups where edge color corresponds to the sender (source) cell subclass and edge thickness reflects the relative interaction strength between cell groups with equivalent maximum edge weight between control and Ts21. (**E**) Violin plots depicting marker gene expression of microglia subtypes Micro *CD200 PRKAA2* (left) and Micro *CTSC DOCK1* (right) in control (green) and Ts21 (purple). (**F**) Transcriptomic UMAP visualization of control microglia using a joint UMAP embedding of both control and Ts21 microglia, colored by control microglia subtype. (**G**) Pie chart depicting the percent representation of control microglia subtypes and total microglia number (below). (**H**) Transcriptomic UMAP visualization of Ts21 microglia using a joint UMAP embedding of both control and Ts21 microglia, colored by control microglia subtype. (**I**) Pie chart depicting the percent representation of Ts21 microglia subtypes and total microglia number (below). (**J and K**) Quantification of the HLA-DRB1 cell total correct fluorescence (CTCF) intensity in control (green; sample age: 2y, 4d postnatal) and Ts21 (purple; sample age: 2y, 57d postnatal) AIF1+ microglia residing in the grey matter (J) or white matter (K). Each dot represents quantification from one tissue section (** p<0.01, **** p<0.0001, two-tailed unpaired t-test). (**L**) Quantification of the HLA-DRB1 cell total correct fluorescence (CTCF) intensity in Ts21 AIF1+ microglia residing in the grey matter (left) or white matter (right). Each dot represents quantification from one tissue section (** p<0.01, two-tailed unpaired t-test).

**Supplementary Fig. 14.**
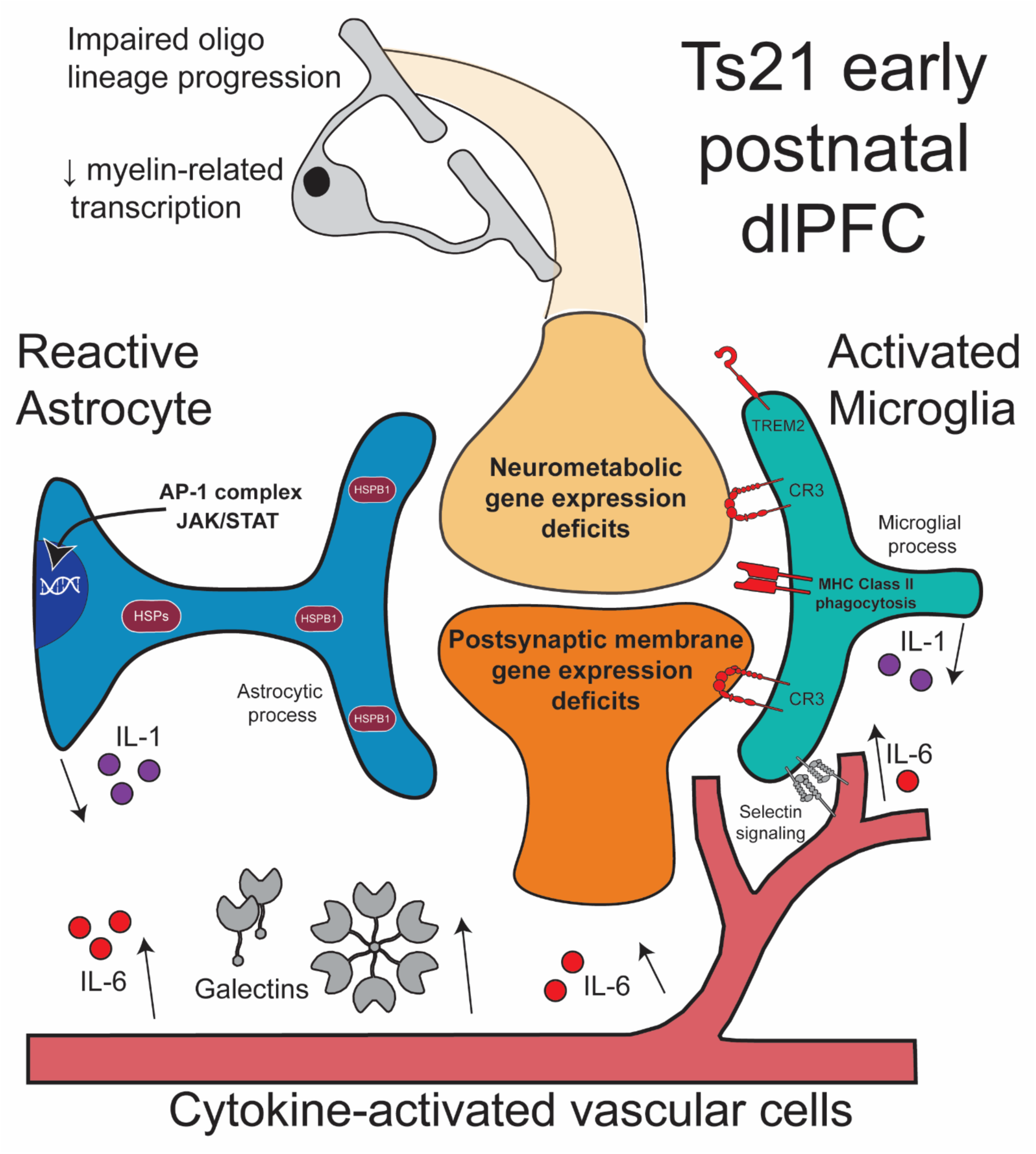
Overview of study findings.

